# Uneven research attention to biodiversity loss revealed by large language models

**DOI:** 10.64898/2026.10.23.761449

**Authors:** Hunain Mohiuddin, Kerstin Forster, Stefan Feuerriegel

## Abstract

Biological diversity is crucial for ecosystem functioning and human well-being but is declining under multiple pressures from human activity. Research on biodiversity loss spans multiple ecological contexts, yet existing syntheses typically cover only subsets of the evidence base. We developed a systematic mapping framework based on large language models to screen and synthesize the scientific literature on biodiversity loss at scale. Applying our framework to 2.4 million bibliographic records, we identified 253,081 scientific articles on biodiversity loss, which we classified across five dimensions: geography, ecosystems, taxa, direct drivers (e.g., land- and sea-use change, climate change, and pollution), and specific threats (e.g., agriculture, hunting, and urban development). Chordates accounted for 34.8% of research attention but only 4.6% of described species. We find comparatively limited research attention to non-chordate animals, studies in Africa, and direct exploitation in marine systems. Overall, our systematic map provides a multidimensional view of research attention to biodiversity loss across various ecological contexts and can help identify evidence gaps to inform future research priorities.

## Main

Biodiversity is declining globally [1–3], with substantial consequences for ecosystem functioning and human well-being [4–6]. The Kunming–Montreal Global Biodiversity Framework sets out the 2050 Vision of “living in harmony with nature” [7] and calls for broad changes across economic, social, political, and technological systems [8]. Addressing biodiversity loss requires a broad understanding of the scientific evidence across the ecological and geographic contexts, including different ecosystems, regions, taxa, and anthropogenic pressures, in which biodiversity change is studied.

However, research on biodiversity change is scattered across different ecological contexts and areas of study. Previous work has documented geographic [9–11] and taxonomic [12–15] differences in research attention, while evidence syntheses have typically focused on specific geographic regions, ecosystems, taxa, direct drivers, or threats [16–21]. In contrast, a comprehensive, multidimensional synthesis of the existing literature across geographic and ecological contexts, taxa, and anthropogenic pressures at scale is still lacking.

Here, we develop a systematic mapping framework that uses large language models (LLMs) to screen and classify the scientific literature on biodiversity change at scale. For this, our framework follows the Collaboration for Environmental Evidence (CEE) guidelines for systematic maps [22]. Our framework first screens bibliographic records for biodiversity change and then codes eligible records along five dimensions: (i) geographic regions, (ii) ecosystems, (iii) taxa, (iv) direct drivers of biodiversity change (e.g., landand sea-use change, climate change, and pollution; defined by the Intergovernmental Science-Policy Platform on Biodiversity and Ecosystem Services [23]), and (v) specific threats (e.g., agriculture, hunting, and urban development; based on the International Union for Conservation of Nature (IUCN) Threats Classification Scheme [24]). We apply our framework to 2.4 million bibliographic records from the Web of Science Core Collection (WoS) published from January 2000 to April 2026. As a result, we assembled a systematic map of 253,081 articles about biodiversity loss. The resulting systematic map supports a global analysis of research attention across geographic regions, ecosystems, taxa, direct drivers, and specific threats, helping to identify areas with limited representation and priorities for future research. The underlying dataset is publicly available at https://osf.io/xg8yq (see Data availability). An interactive dashboard for exploring the systematic map is available at https://biodiversityliterature.com/.

**Figure 1:**
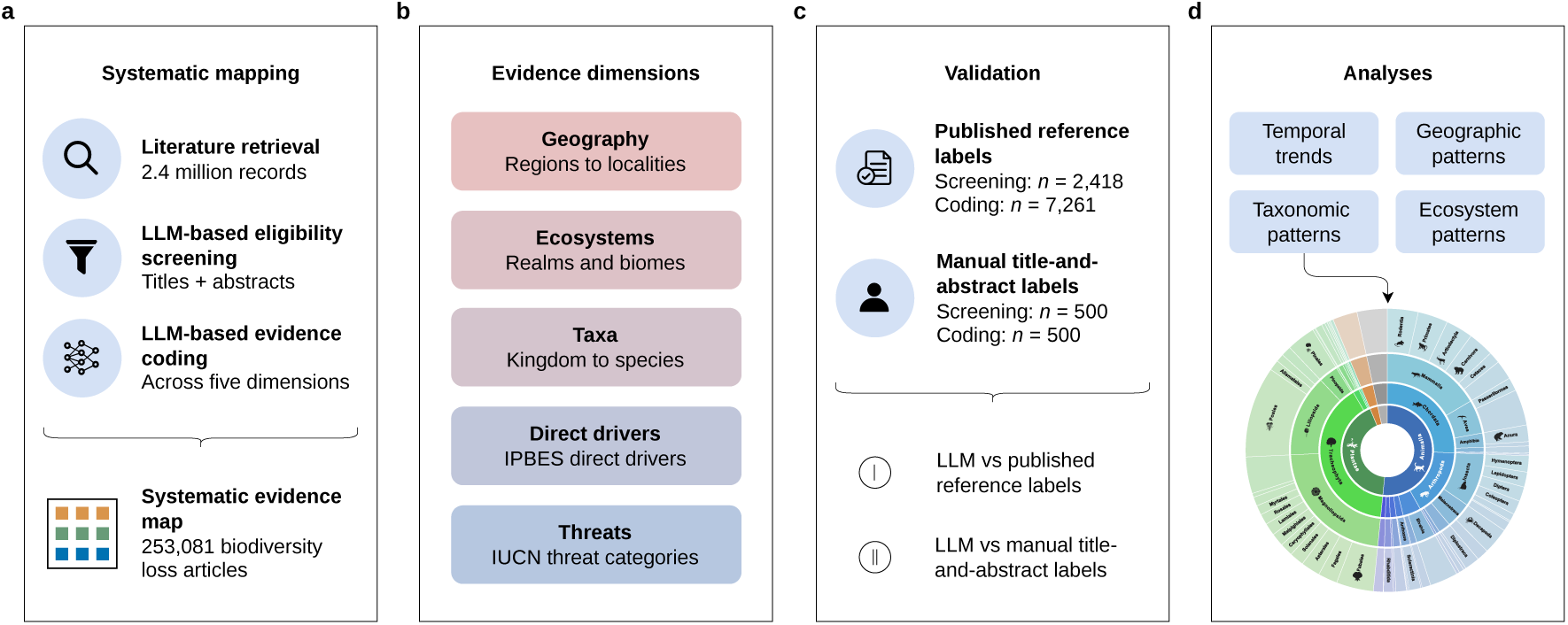
LLM-based systematic mapping of scientific evidence on biodiversity loss. (**a**) The systematic mapping workflow comprised three steps: (1) literature retrieval from the Web of Science Core Collection, (2) eligibility screening based on titles and abstracts, and (3) multidimensional evidence coding. The resulting evidence map covered 253,081 scientific articles on biodiversity loss. (**b**) We used five dimensions for evidence coding: geography, ecosystems, taxa, direct drivers (defined by the Intergovernmental SciencePolicy Platform on Biodiversity and Ecosystem Services; IPBES [23]), and threats (based on the International Union for Conservation of Nature (IUCN) Threats Classification Scheme [24]). (**c**) Screening and coding were validated against (i) published reference labels and (ii) manual title-and-abstract annotations (Supplementary Methods S3). (**d**) The resulting evidence map was analyzed across time, geography, taxa, and ecosystems.

## Results

### Research on biodiversity loss is rapidly expanding

We screened 2.4 million bibliographic records and identified 253,081 scientific articles with a thematic focus on biodiversity loss. We classified these scientific articles across five dimensions: (1) the geographic regions defined by the Intergovernmental SciencePolicy Platform on Biodiversity and Ecosystem Services (IPBES) [25], (2) the ecosystems according to the Global Ecosystem Typology [26], (3) the taxa according to the Global Biodiversity Information Facility (GBIF) Backbone Taxonomy [27], (4) direct drivers defined by the IPBES [23], and (5) threats according to the International Union for Conservation of Nature (IUCN) Threats Classification Scheme [24]. In the following, we use “research attention” to refer to the share of mapped scientific articles and thus as a proxy for the representation of different research themes in the literature. All analyses are based on complete publication years from 2000 to 2025 (*n* = 245,168). Unless stated otherwise, we report research attention among articles with a resolved classification for the respective dimension; where an article was assigned to multiple categories within a dimension, the contribution was divided proportionally across categories.

The annual number of mapped scientific articles on biodiversity loss increased more than eightfold, from 2,530 in 2000 to 20,520 in 2025 (see Figure 2a). The composition of research attention across the broader threat categories changed over the same period. The share assigned to climate change and severe weather increased from 4.9% to 19.5%, while pollution remained the largest category but declined from 40.5% to 32.5% (Figure 2b; Supplementary Table S1).

**Figure 2:**
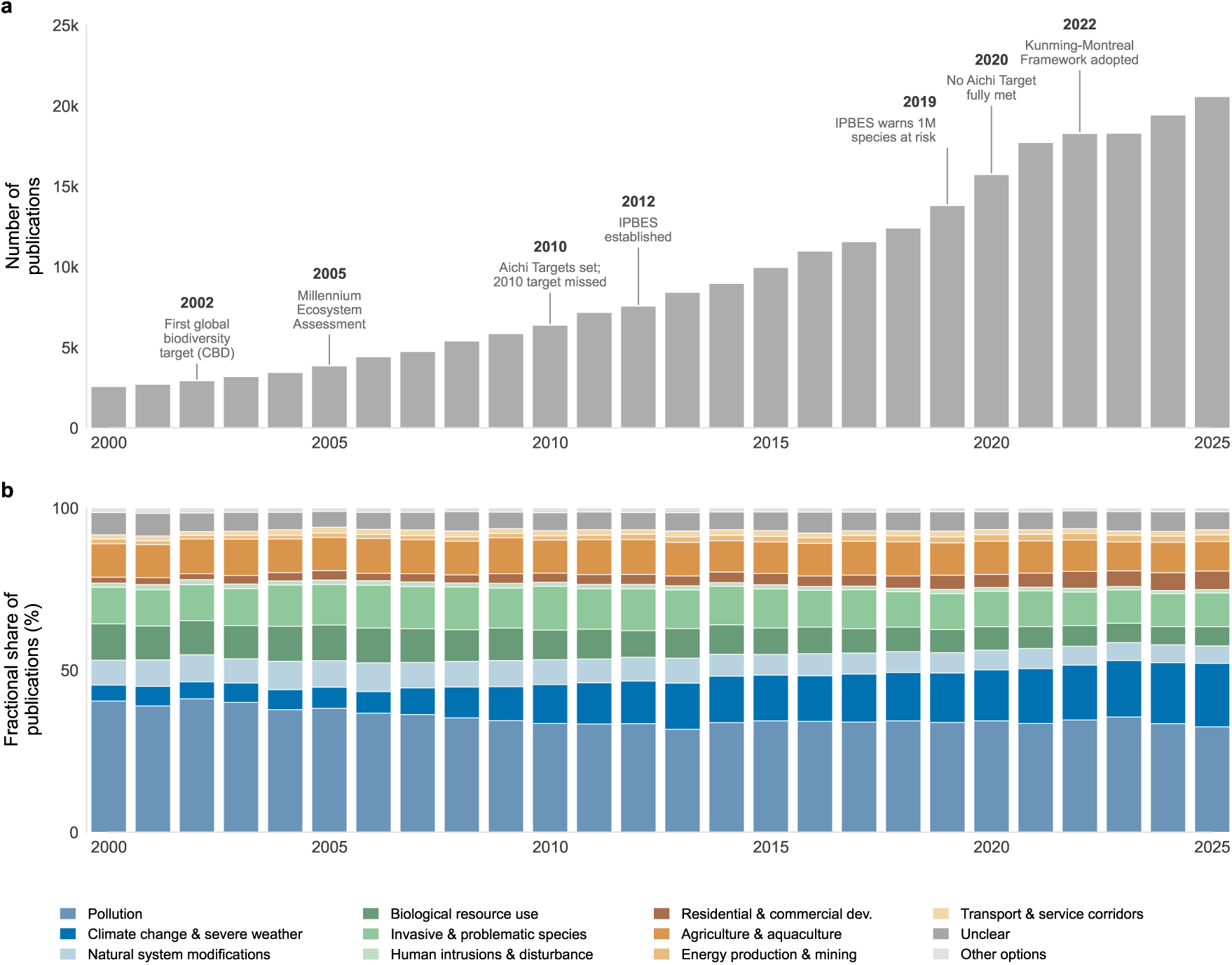
Temporal development of the scientific evidence on biodiversity loss. (**a**) Annual number of scientific articles identified as reporting biodiversity loss from 2000 to 2025 (*n* = 245,168). The annotations show selected international biodiversity policy and assessment milestones. (**b**) Annual distribution of research attention across broad threat categories from the International Union for Conservation of Nature (IUCN) Threats Classification Scheme [24], where threats describe specific anthropogenic pressures on biodiversity (e.g., agriculture, pollution, or biological resource use).

### Research attention to biodiversity loss is uneven across geographies

Research attention was concentrated in a small number of countries (Figure 3a), consistent with previous evidence of geographic imbalances in biodiversity research [9, 10]. The United States accounted for 15.5% of article–country attributions, followed by China (12.5%) and Brazil (5.2%); together, the three countries accounted for 33.2%. In contrast, 192 of the 213 countries and territories each accounted for less than 1% of these attributions (Supplementary Figure S4).

**Figure 3:**
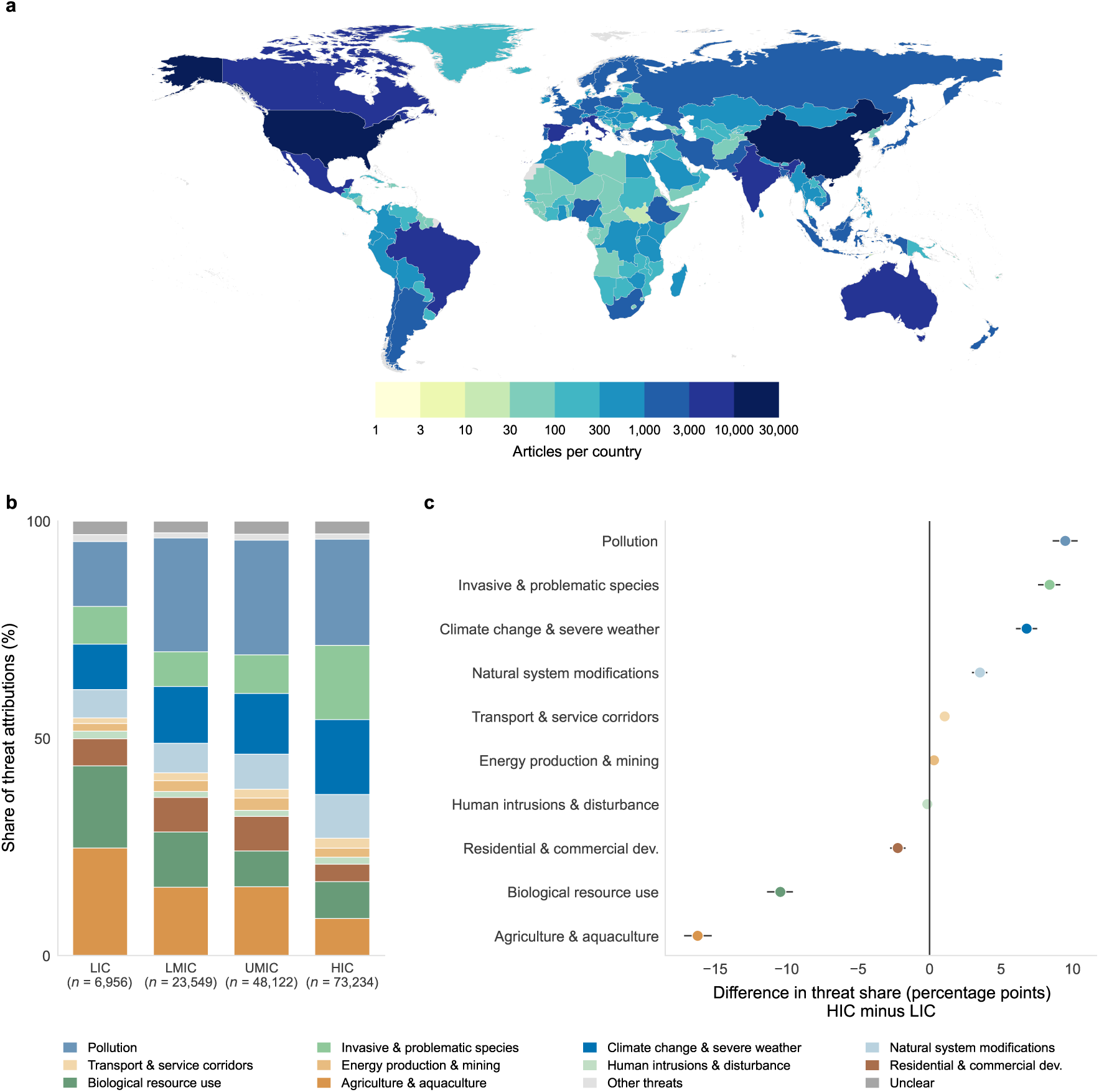
Geographic distribution of research attention to biodiversity loss. (**a**) Geographic distribution of articles linked to at least one study country (*n* = 146,630). (**b**) Research attention by World Bank income group [28]: low-income (LIC), lower-middleincome (LMIC), upper-middle-income (UMIC), and high-income (HIC). Each bar is further divided by broad IUCN threat categories [24]. For threat shares, each article contributes a total weight of one, divided equally among its study countries and assigned threat categories. (**c**) Difference in threat shares between HIC and LIC (HIC minus LIC, percentage points). Positive values indicate higher shares in HIC countries and negative values a higher share in LIC countries. Points show calculated differences; whiskers show 95% bootstrap confidence intervals.

To examine how research attention differs across economic contexts, we grouped the countries subject to study according to the income groups defined by the World Bank [28]. Among article–income-group attributions, high-income countries (HIC) accounted for 48.2% (*n* = 73,234), followed by upper-middle-income countries (UMIC) with 31.7% (*n* = 48,122), lower-middle-income countries (LMIC) with 15.5% (*n* = 23,549) and low-income countries (LIC), 4.6% (*n* = 6,956). We further compared the composition of the broad IUCN threat categories (i.e., which capture specific anthropogenic pressures on biodiversity) across the income groups (see Figure 3b). For each threat category, we calculated the difference in the share of research attention between HIC and LIC evidence (i.e., HIC minus LIC; Figure 3c). Agriculture and aquaculture received a larger share of research attention for LIC than HIC countries, with a difference of 16.2 percentage points, followed by biological resource use (10.4 percentage points). In contrast, pollution received a larger share of research attention in HIC countries (9.5 percentage points), followed by invasive and problematic species (8.4), climate change and severe weather (6.8), and natural system modifications (3.5).

### Research attention is concentrated on a subset of taxa

To examine research attention across taxa, we matched taxonomic information from the mapped articles to the GBIF Backbone Taxonomy [27]. Taxa could be identified for 171,791 articles (70.1%). For comparisons across the taxonomic hierarchy, we retained taxonomic paths represented in both the mapped evidence and the GBIF benchmark, yielding 113,342 articles at the order level.

Research attention varied substantially across the taxonomic hierarchy (Figure 4). At the kingdom level (innermost ring), Plantae accounted for 42.2% of research attention and 17.1% of described species. At the phylum level (second ring), Chordata accounted for 25.6% and 4.6%, respectively, compared with 16.2% and 51.5% for Arthropoda. At the class level (third ring), Mammalia accounted for 16.5% of research attention and 0.81% of described species, compared with 9.0% and 42.4% for Insecta. A UMAP projection visualizes similarities among articles across taxonomic classes (Supplementary Figure S9). To compare general patterns, we grouped taxa into five categories (chordates, nonchordate animals, plants, fungi, and other taxa), which we refer to as broader categories below. Within this grouping, chordates accounted for 34.8% of research attention but only 4.6% of described species (Supplementary Figure S6). Research attention to both chordates and non-chordate animals declined over time. For chordates, research attention remained comparatively limited in Africa relative to the broader distribution of threatened-species occurrences reported by the IUCN [29], while its geographic distribution shifted toward Asia and the Pacific.

**Figure 4:**
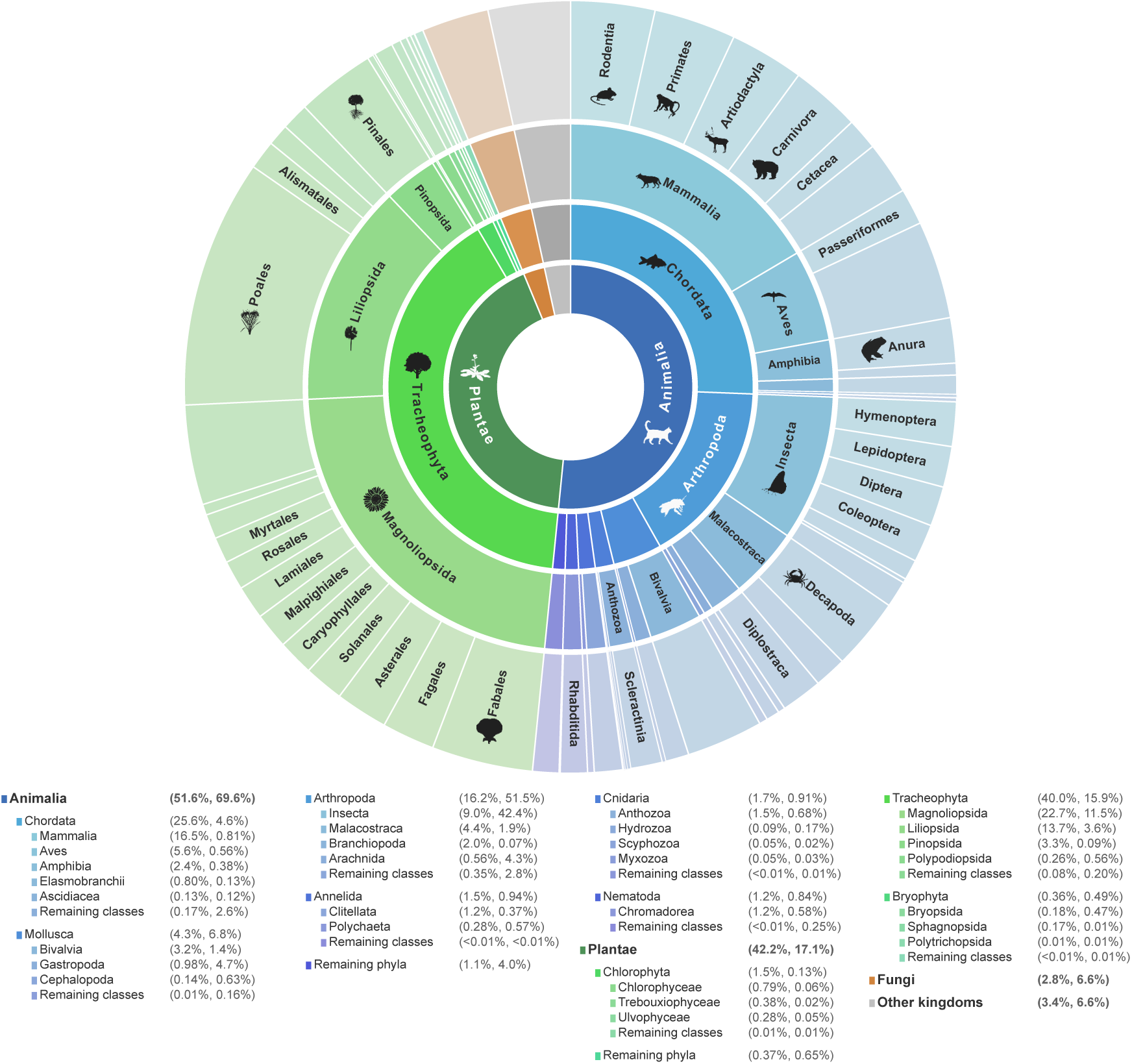
Research attention across taxa. Reported taxa were matched to the GBIF Backbone Taxonomy [27]. From the center outward, rings represent kingdom, phylum, class, and order, with segment size proportional to research attention. For each listed taxon, percentages indicate its share of research attention in the matched article subset and its share of accepted species in the fixed GBIF benchmark (*n* = 2,607,214), respectively. The matched article subset varies across taxonomic ranks, with lower coverage for chordates partly reflecting differences in fish and reptile classifications. Taxa not displayed individually are aggregated as “Remaining” where indicated; Fungi and smaller kingdoms are shown only at kingdom level.

### Research attention to direct drivers varies across terrestrial, freshwater, and marine systems

To compare research attention across terrestrial, freshwater, and marine systems, we examined the five direct drivers defined by IPBES [30, 31], which capture major anthropogenic pressures that directly affect biodiversity: land- and sea-use change, direct exploitation, climate change, pollution, and invasive alien species (see Figure 5). Previously, global assessments identified land/sea use change as the leading direct driver in terrestrial and freshwater systems and direct exploitation in marine systems [30, 31]. In our evidence map, land/sea use change received the largest share of research attention in terrestrial systems (32.7%), whereas pollution accounted for the largest share in both freshwater (57.4%) and marine systems (38.3%). Direct exploitation accounted for 10.5%, 6.7%, and 24.5% of research attention in terrestrial, freshwater, and marine systems, respectively, and was not the leading driver in any of them. Research attention therefore showed a similar pattern to global assessments in terrestrial systems, while the distribution differed in freshwater and marine systems. In freshwater systems, the prominence of pollution occurred alongside other pressures highlighted in global assessments, including flow alteration and habitat degradation [32]. UMAP projections visualize similarities among articles across ecosystems (Supplementary Figure S7) and direct drivers (Supplementary Figure S8).

**Figure 5:**
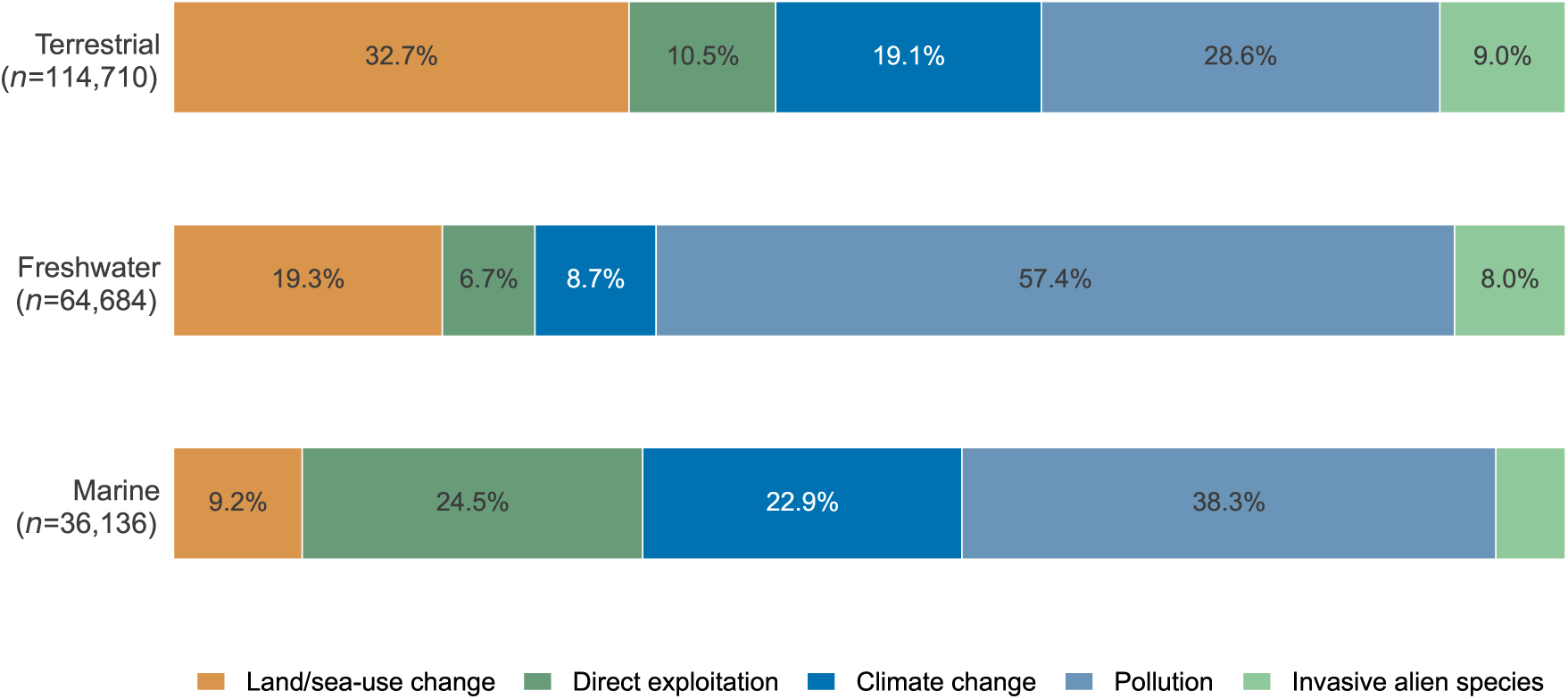
Distribution of research attention across direct drivers in terrestrial, freshwater, and marine systems. Bars show the share of research attention across the five direct drivers defined by the Intergovernmental Science-Policy Platform on Biodiversity and Ecosystem Services (IPBES) [23], which capture major anthropogenic pressures that directly affect biodiversity. Results are shown separately for terrestrial (*n* = 114,710), freshwater (*n* = 64,684), and marine (*n* = 36,136) systems. Articles linked to multiple drivers contribute equally across them, so that shares sum to 100% within each system. Only articles assigned to exactly one of the three systems were included.

Research attention across drivers also changed over time. The share assigned to direct exploitation declined in all three systems, while pollution declined in terrestrial research and showed an initial decline followed by a partial increase in freshwater and marine research. Within pollution-related articles, mentions of plastics or marine debris increased substantially over the study period, reaching 9.6% in terrestrial, 12.4% in freshwater, and 24.0% in marine research by 2025 (see Supplementary Figure S5).

## Discussion

We developed an LLM-based systematic mapping framework to screen 2.4 million bibliographic records and identified 253,081 scientific articles focused on biodiversity loss. Our evidence map shows that research on biodiversity loss has expanded substantially, but research attention remains uneven across geography, ecosystems, taxa, direct drivers, and threats. Existing syntheses based on manual screening are typically limited to subsets of taxa [33–37], geographies [17, 38–41], direct drivers [19, 20, 34, 42–44], or ecosystems [41, 44–46]. Our framework, in contrast, offers a comprehensive, multidimensional view of research attention across these dimensions at scale within a single evidence map. The growth in the number of scientific publications on biodiversity loss is consistent with the broader expansion of biodiversity science and has occurred alongside increasing institutional and policy attention [47–52]. At the same time, research attention remains uneven across countries. In particular, high-income countries accounted for almost half of all article–income-group attributions. These patterns may reflect differences in scientific capacity, consistent with previous studies showing substantial cross-country variation in biodiversity and ecological research [9, 10, 53], and reflect heterogeneity in species richness, endemism, and extinction risk [54–56]. For example, Brazil, which encompasses the Atlantic Forest and Cerrado, two recognized global biodiversity hotspots [55], ranks third in terms of research attention in our evidence map. In contrast, Africa received comparatively little research attention, consistent with previous studies on regional differences in research coverage and scientific capacity [9, 10].

Research attention also differs substantially across taxa. For example, the geographic distribution of chordate research shows large regional differences. Previous studies have reported similar taxonomic imbalances in biodiversity research [9, 11, 12, 14] and species occurrence data [13], while, in contrast, our work provides a systematic and multidimensional assessment of such differences across biodiversity loss research at scale. Comparing research attention with described species diversity can further help identify taxa that are potentially underrepresented in the literature. Similar patterns are also visible for specific anthropogenic pressures on biodiversity. For example, agriculture and biological resource use received greater research attention in low-income countries, broadly consistent with global assessments identifying agriculture, logging, and hunting as major threats across many tropical regions [57, 58]. Across ecosystems, pollution received the most research attention in freshwater and marine systems, contrasting with global assessments that identify land- and sea-use change and direct exploitation, respectively, as the leading drivers [30, 31]. Together, these patterns highlight areas with comparatively limited representation in the literature.

Our LLM-based framework has several strengths. First, it provides a comprehensive, multidimensional mapping at a scale that would be difficult to achieve through manual screening and coding alone. Overall, our framework applies a consistent classification across our evidence base of 253,081 articles reporting biodiversity loss. Second, the use of a common classification framework allows research attention to be compared consistently across geography, ecosystems, taxa, direct drivers, and specific threats. Third, our framework is designed to be updated as the scientific literature grows (similar to “living reviews” [59]). We thus make our framework and dataset publicly available to support reproducibility and future updates.

Our evidence map should be interpreted in light of several limitations. First, we restricted our search to scientific articles indexed in Web of Science, thereby excluding gray literature. Second, screening and coding were based on titles and abstracts rather than full texts. This is consistent with previous large-scale approaches that use automated text analysis to map scientific literature in sustainability research [60–64]. Third, the classifications were generated by an LLM, which itself may introduce errors but allow scientific articles to be screened and classified consistently at scale. We therefore validated our framework against published reference labels and manual title-and-abstract annotations (see Supplementary Methods S3). The overall performance of evidence coding was strong, but varied across classification fields, and classifications tended to be more reliable at broader categorizations than for granular classifications such as species-level identities. Eligibility screening prioritized recall over precision. Fourth, not all articles contained sufficient information to assign a classification in every dimension. Individual analyses therefore use the subset of articles for which the relevant dimension could be resolved. Fifth, the evidence map describes patterns of research attention in the scientific literature and does not quantify ecological effect sizes or the relative importance of different pressures.

In sum, by tracking how research attention is distributed across the biodiversity literature, our evidence map can help identify both evidence gaps and ecological contexts that remain sparsely represented. This can inform future research agendas, target scientific efforts, and support a systematic use of existing evidence.

## Methods

We developed an LLM-based framework for systematically mapping the research on biodiversity loss. Our study followed the Collaboration for Environmental Evidence (CEE) guidance for systematic maps [22], which we adapted for LLM-based screening and coding, and was preregistered [65]. We provide a ROSES flow diagram to show the search process in Supplementary Figure S2.

### LLM-based framework for mapping biodiversity loss research

Our LLM-based framework comprises three steps: (1) literature retrieval, (2) eligibility screening, and (3) evidence coding (Figure 1). Evidence coding followed predefined criteria derived from established biodiversity classification frameworks for geography, ecosystems, taxa, direct drivers, and threats [24–27, 66]. We validated the screening and coding steps against published reference labels and manual annotations (see Supplementary Methods S3). The overall design of our LLM framework was informed by previous studies using machine learning for large-scale bibliographic analyses in environmental and sustainability research [60–64, 67]. Our reporting follows the GUIDE-LLM checklist [68].

**Step 1: literature retrieval.** We searched the Web of Science Core Collection (WoS) for scientific articles published between 1 January 2000 and 30 April 2026 without language restrictions. To prioritize recall during retrieval, we combined biodiversity-relevant WoS subject categories with terms describing biodiversity change or assessment and term blocks covering the five IPBES direct drivers [31] (see Supplementary Table S2). To focus on research published in indexed scientific journals, we restricted our search to articles in the WoS, thereby excluding gray literature. We finally removed duplicate records as well as records without an abstract. In total, around 2.4 million records entered the subsequent eligibility screening.

We evaluated the search against an a priori list of 581 WoS-indexed articles drawn from five published sources [17, 19–21, 31] and the direct-driver chapters of the IPBES Global Assessment [23]. Of the 537 articles published within the search window (2000 onwards), the search retrieved 464, corresponding to a recall of 86.41% (Supplementary Table S3).

**Step 2: eligibility screening.** We screened records using GPT-5-nano (version `2025-08-07`) accessed through the OpenAI API. We selected GPT-5-nano for its strong performance in text classification tasks and its comparatively low computational cost, which is necessary to screen millions of bibliographic records. Screening was based on the title and abstract of each record, which is consistent with previous large-scale LLM-based bibliographic analyses [60–64, 67].

Informed by best practices in prompt engineering [69–71], we used a schemaconstrained prompt designed to capture different ways in which articles report biodiversity change and its relationship to anthropogenic pressures. Specifically, the prompt distinguished between two aspects: (i) whether biodiversity change was reported and its direction, and/or (ii) whether direct anthropogenic drivers were reported or plausibly implied and linked to that change (see Supplementary Figure S1). We used this multi-dimensional approach because articles may report biodiversity change without specifying its direction or drivers, or discuss anthropogenic pressures without explicitly linking them to biodiversity change. We operationalized these aspects through four screening questions (i.e., whether the article reported biodiversity change, the direction of that change, whether a direct anthropogenic driver was reported or plausibly implied, and whether the driver was linked to the biodiversity change), each returning `Yes`, `Unclear`, or `No`. To prioritize re-call, we retained records unless at least one question returned `No`, allowing for incomplete information in titles and abstracts. Only 123 records received an `Unclear` classification for all four screening questions; we retained all of them after confirming through manual inspection that this subset was generally relevant and fitted our eligible criteria.

We subsequently identified articles reporting negative biodiversity change for the analysis of biodiversity loss. We validated the eligibility screening against human assessments (see below), finding that the procedure achieved high recall. The prompt, output schema, and model configuration are described in Supplementary Methods S5.1.

Applying this approach to the full corpus, we identified 605,199 eligible records. Of these, 253,081 articles were classified as reporting biodiversity loss and formed the evidence map analyzed in this study (Supplementary Methods S2).

**Step 3: evidence coding.** We classified the title and abstract of each eligible record across five dimensions: (1) geographic regions based on the IPBES geographic classification [25], (2) ecosystems based on the Global Ecosystem Typology [26], (3) taxa based on the GBIF Backbone Taxonomy [27], (4) direct anthropogenic drivers based on the five IPBES direct drivers [23], and (5) specific threats based on the IUCN Threats Classification Scheme [24]. For the classification, we again used GPT-5-nano (version `2025-08-07`), accessed through the OpenAI API, with a separate schema-constrained prompt for each dimension (see Supplementary Methods S5.2 for prompts, output schemas, and model configuration). Within each dimension, an article could receive several labels (Supplementary Table S10). We standardized the geographic identifiers in these outputs and matched the taxon names to the GBIF Backbone Taxonomy to obtain article-level labels (Supplementary Methods S4.1 and S4.2).

### Validation

We validated LLM-based eligibility screening and evidence coding against two complementary sources of validation data: (i) published reference labels derived from full-text assessments in previous evidence syntheses, and (ii) manual title-and-abstract annotations (Supplementary Methods S3.1). Both sources were used to evaluate LLM-based eligibility screening (Supplementary Methods S3.2) and evidence coding (Supplementary Methods S3.3).

Against published reference labels, LLM-based screening achieved a recall of 90.5% on the held-out test set (*n* = 482). For evidence coding, the LLM achieved a macro-averaged weighted precision of 85.10%, recall of 82.79%, F1 of 83.33%, and Cohen’s *κ* of 0.74 across five extracted dimensions. Performance varied across classification fields, with F1 scores of 90.38% for geographic regions, 92.61% for ecosystems, 83.85% for direct drivers, 68.43% for broad IUCN threats, and 57.95% for species-level classifications (Supplementary Table S7).

Against manual title-and-abstract labels, LLM screening achieved a precision of 64.5%, recall of 90.1%, and F1 of 75.2% on a sample of 400 records drawn directly from the search results. For evidence coding, we compared LLM classifications with 500 dimension-specific manual annotations (100 per dimension), covering 479 distinct articles. The LLM achieved a macro-averaged weighted precision of 87.48%, recall of 87.94%, F1 of 86.74%, and Cohen’s *κ* of 0.82 compared to the manual labels (Supplementary Table S8).

### Analysis

Our evidence map on biodiversity loss comprised 253,081 articles. We restricted the primary analyses to the 245,168 articles published from 2000 to 2025 to ensure that all years included in the temporal comparisons were complete. To examine temporal development, we counted mapped articles by publication year and calculated the annual distribution of research attention across broad IUCN threat categories. To calculate annual shares of research attention across broad IUCN threat categories, we divided each article contribution equally among the assigned categories, so that each article contributed a total weight of one. For geographic analyses, we counted each article once per study country in the country map, while compositional research attention shares divided each article contribution equally among its assigned countries. Countries were identified using standardized ISO3 codes and assigned to World Bank income groups according to their classification in the article publication year [28]. For each broad IUCN threat category, we calculated the difference in its share of research attention between high-income and low-income countries (HIC minus LIC) and estimated 95% bootstrap confidence intervals by resampling articles. For taxonomic analyses, we matched coded taxa to the GBIF Backbone Taxonomy [27] and compared their representation in the mapped literature with accepted described species in a fixed GBIF Backbone snapshot. We examined five broad groups (chordates, nonchordate animals, plants, fungi, and other taxa) as well as taxonomic classifications at the kingdom, phylum, class, and order levels. For chordates, we additionally compared the regional distribution of research attention across the four IPBES regions with the regional distribution of threatened-species occurrences in the IUCN benchmark. To compare direct drivers across ecosystems, we used the five IPBES direct drivers, which allow comparison with global assessments of driver importance, and restricted the analysis to articles assigned to exactly one of the three ecosystems (i.e., terrestrial, freshwater, or marine). Further details on the geographic, taxonomic, and ecosystem analyses are provided in Supplementary Methods S4.

## Data availability

The processed data generated in this study are available via https://osf.io/xg8yq/overview. WoS-derived bibliographic metadata cannot be redistributed under Clarivate’s end-user terms. Instead, our repository provides record identifiers and instructions for reproducing the metadata. An interactive dashboard for exploring the evidence map is available at https://biodiversityliterature.com/.

## Code availability

The code used for the LLM-based screening, evidence coding, and all analyses reported in this study is available on GitHub at https://github.com/forsterkerstin/biodiversity-loss-systematic-map. The repository also contains the prompts, configuration files, and instructions for reproducing the workflow.

## Author contributions

H.M. designed the study, developed the LLM-based screening and coding pipeline, performed the analysis, and wrote the manuscript. K.F. and S.F. edited the manuscript and provided feedback. S.F. conceived the study. All authors read and approved the final manuscript.

## Funding

No external funding was received.

## Competing interests

The authors declare no competing interests.

## Supplementary Figures

**Supplementary Figure S1:**
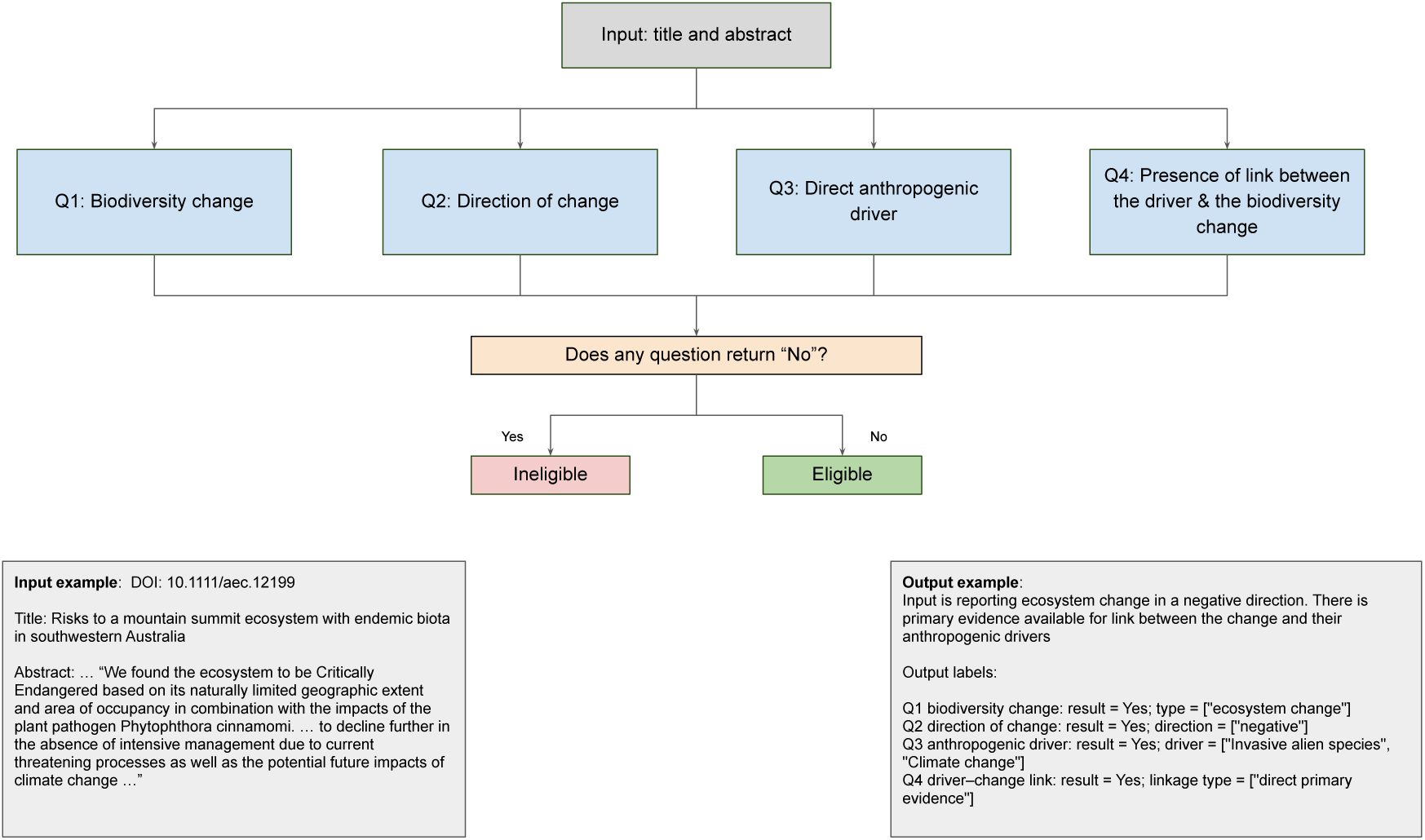
Eligibility screening procedure. Each record is screened using four questions (`Q1`–`Q4`) based on the title and abstract: `Q1` biodiversity change (does the record report at least one form of biodiversity change, operationalized across genetic, species, community, and ecosystem change?); `Q2` direction of change (does the record identify a direction of biodiversity change, recorded as negative, positive, mixed, or unclear? Here, a negative change, i.e., loss, is the primary analytical focus of the Results); `Q3` direct anthropogenic driver (does the record mention or imply at least one driver, assessed using the five IPBES driver categories and the IUCN Threats Classification Scheme?); and `Q4` linkage (does the record report or imply a link between the identified driver and the biodiversity change?). A record is classified as eligible only when all four questions return `Unclear` or `Yes`; a single `No` at any question yields an ineligible classification. Each question produces a structured label alongside the results. The full operationalization is provided in the LLM prompt in the code.

**Supplementary Figure S2:**
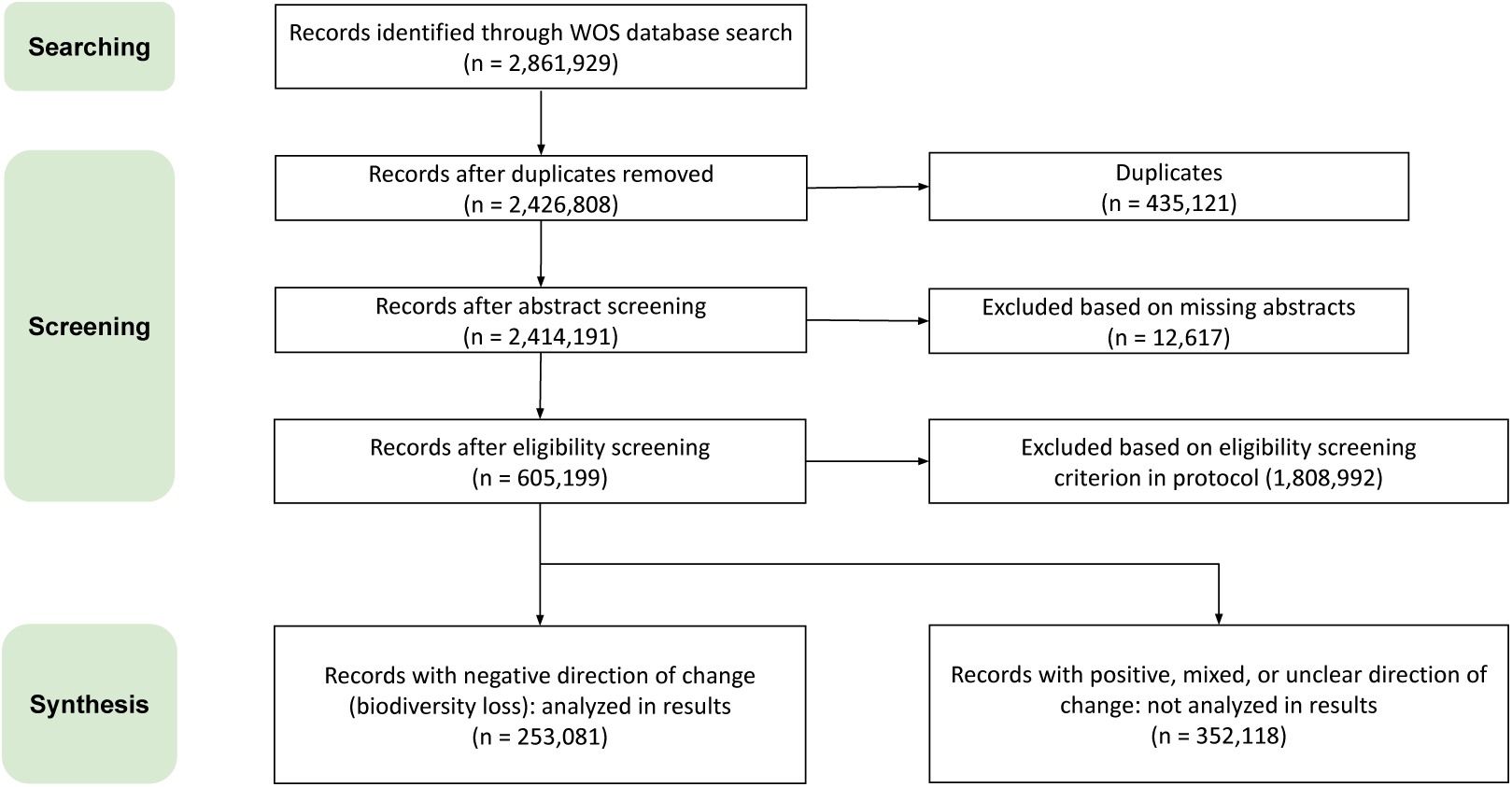
ROSES flow diagram. Of the 605,199 records meeting the pre-registered eligibility criteria, 253,081 articles were classified as reporting biodiversity loss and formed the evidence map analyzed in this study. The remaining 352,118 eligible records, classified as reporting positive, mixed, or unclear biodiversity change, were retained in the released dataset but are not the focus of the present analyses. A detailed breakdown of screening exclusions is provided in Supplementary Figure S3.

**Supplementary Figure S3:**
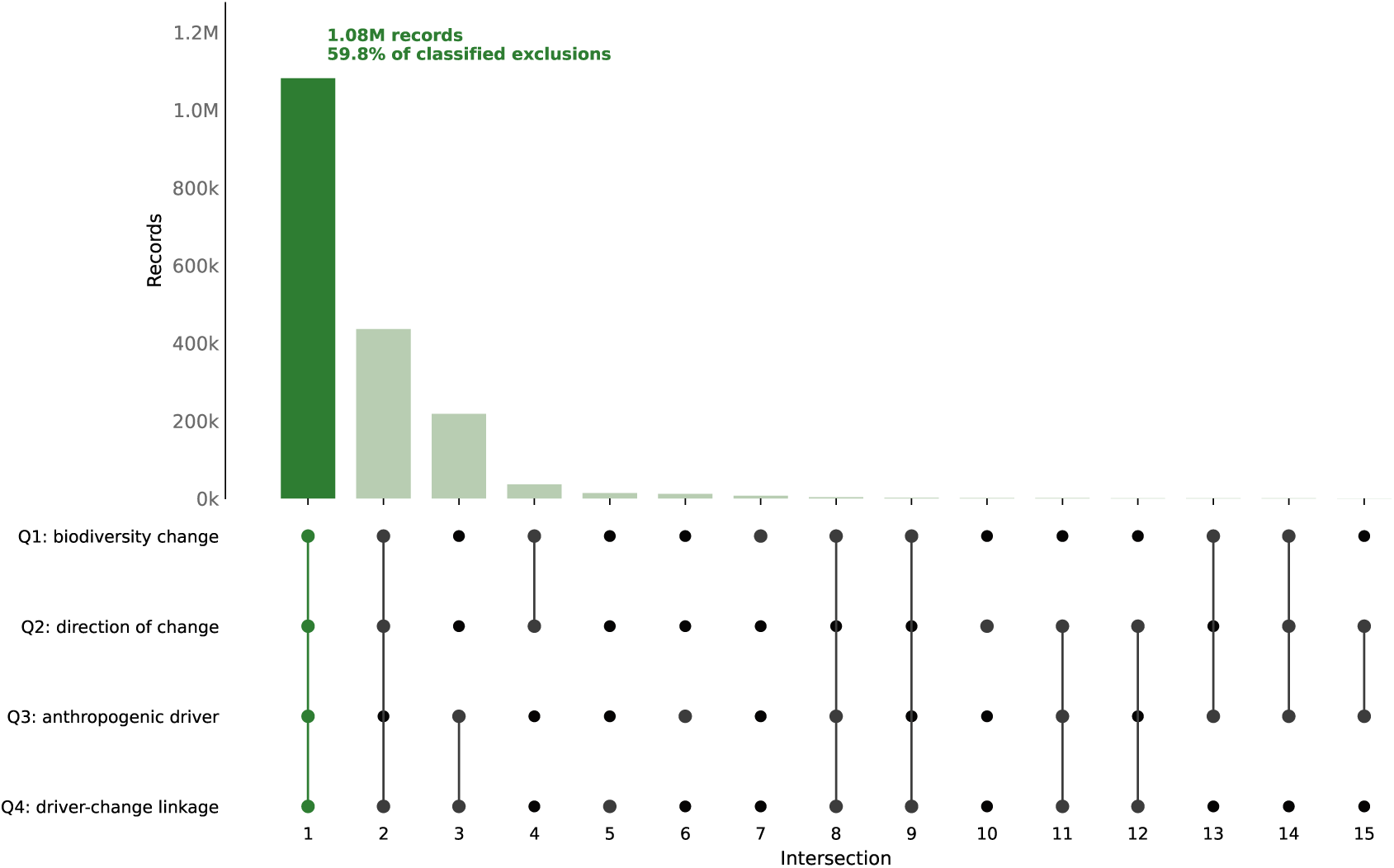
Breakdown of excluded records by screening criteria. Each column shows a different combination of screening questions (`Q1`–`Q4`; see Supplementary Figure S1). For each combination, the corresponding number of excluded records is shown in the bar chart (i.e., when the corresponding questions are answered all with `No`). Most exclusions failed all four criteria simultaneously (*n* = 1,081,249, 59.8% of classified exclusions).

**Supplementary Figure S4:**
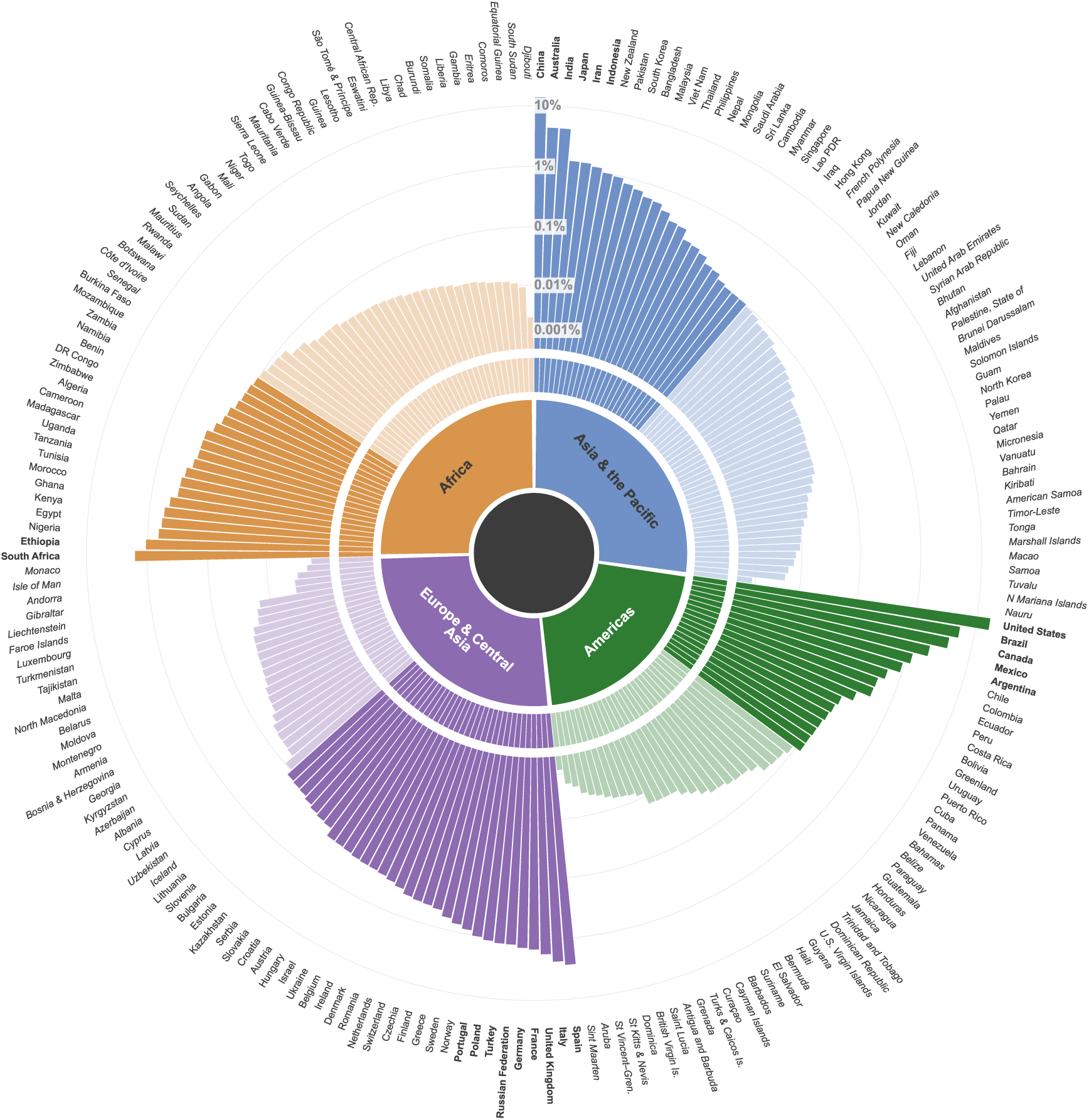
Geographic distribution of research attention across countries. The figure shows the relative distribution of research attention *per* geographic region. Countries and territories are grouped by IPBES region and ordered clockwise by decreasing research attention. Percentages show each country’s share of research attention, with each article contribution divided equally among the study countries. Country names are formatted bold for shares above 1%; countries with a share below 0.1% are shown in a pale color.

**Supplementary Figure S5:**
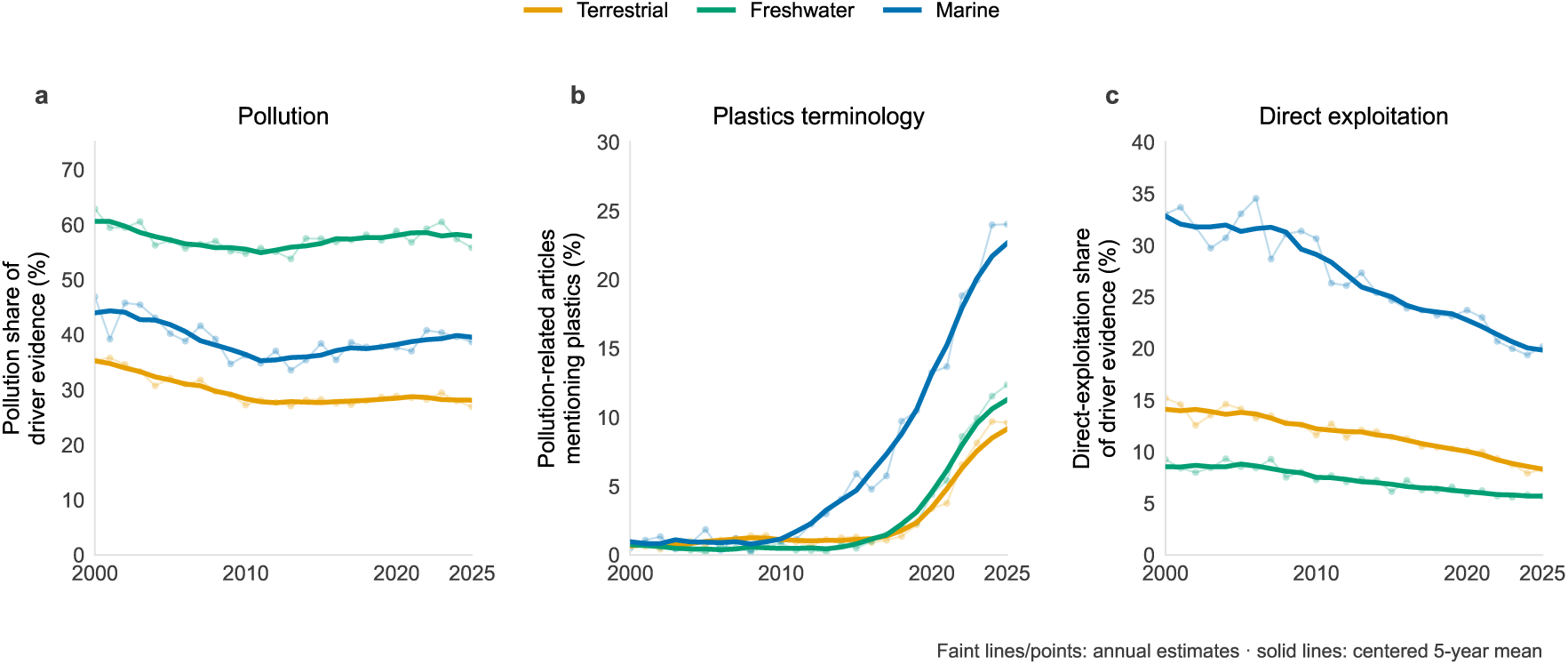
Changes in research attention to pollution and direct exploitation across terrestrial, freshwater, and marine systems, 2000–2025. Thin lines show annual estimates and thick lines show centered five-year moving averages. (**a**) Share of direct-driver research attention assigned to pollution. (**b**) Percentage of pollution-coded articles containing plastics- or debris-related terminology in their title or abstract. Terms were identified using case-insensitive searches for “plastic(s)”, “microplastic(s)”, “nanoplastic(s)”, combinations of “plastic” with “debris”, “waste”, “litter”, “particle(s)”, “fibre(s)/fiber(s)”, “pellet(s)”, “fragment(s)”, or “pollution”, as well as “marine debris/litter” and “anthropogenic litter”. Word boundaries were used to avoid unrelated matches. Percentages were calculated among pollution-coded articles with available title or abstract text. (**c**) Share of direct-driver research attention assigned to direct exploitation. Panels (**a**) and (**c**) use proportionally allocated driver shares, whereas panel (**b**) reports the unweighted percentage of pollution-coded articles containing the specified terminology.

**Supplementary Figure S6:**
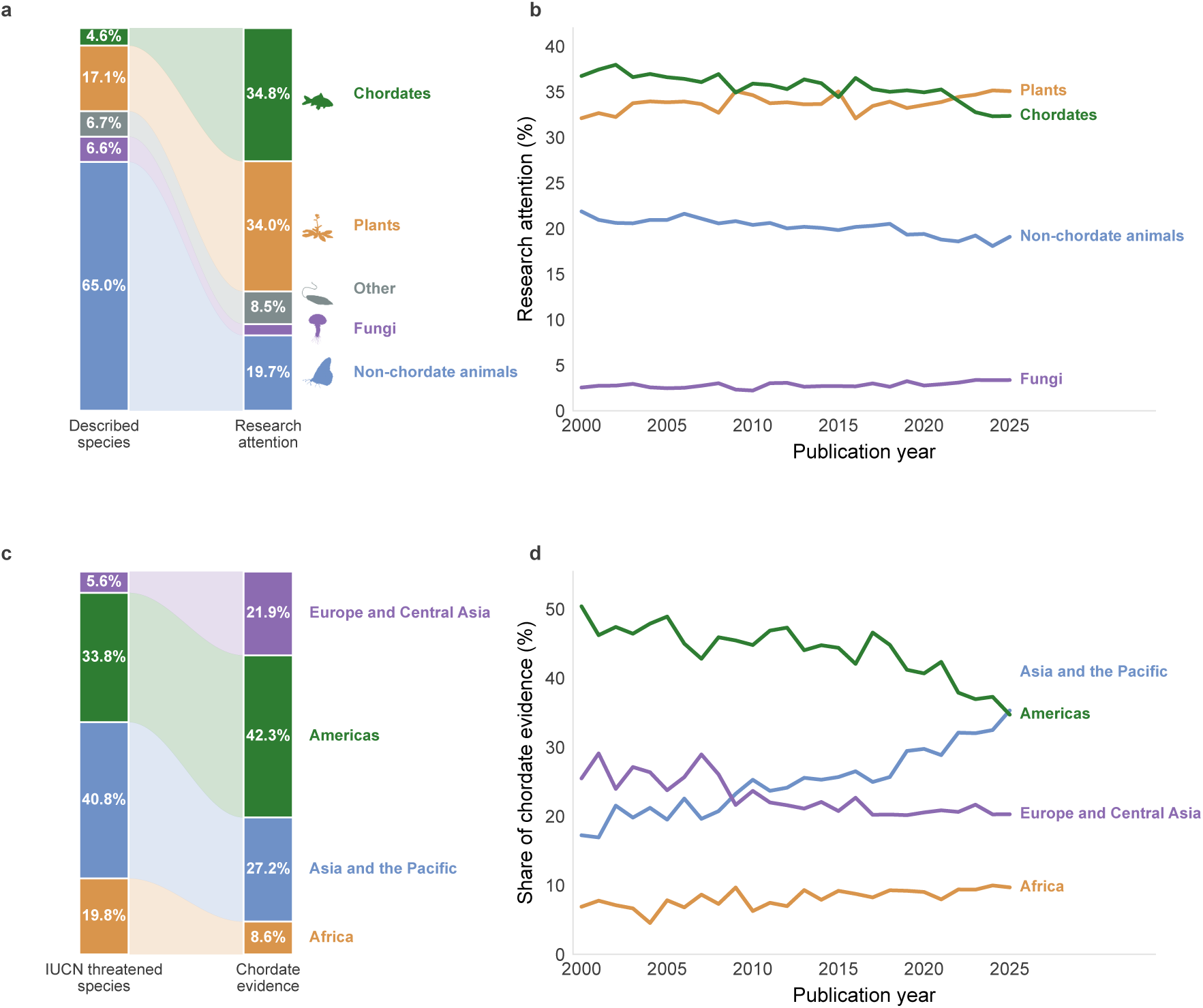
Research attention compared with counts for described species and threatened species. (**a**) Research attention across taxonomic groups compared with their share of described species. Described species are based on the Global Biodiversity Information Facility Backbone Taxonomy. (**b**) Changes in research attention across taxonomic groups over time. (**c**) Regional distribution of chordate research attention compared with the distribution of IUCN Red List threatened-species occurrences (Critically Endangered, Endangered, or Vulnerable) aggregated across IPBES subregions [29]. The reference includes all comprehensively assessed taxonomic groups rather than chordates alone. (**d**) Changes in the geographic distribution of chordate research over time. Panels (**a**) and (**c**) cover 2000–2025; panels (**b**) and (**d**) show annual values.

**Supplementary Figure S7:**
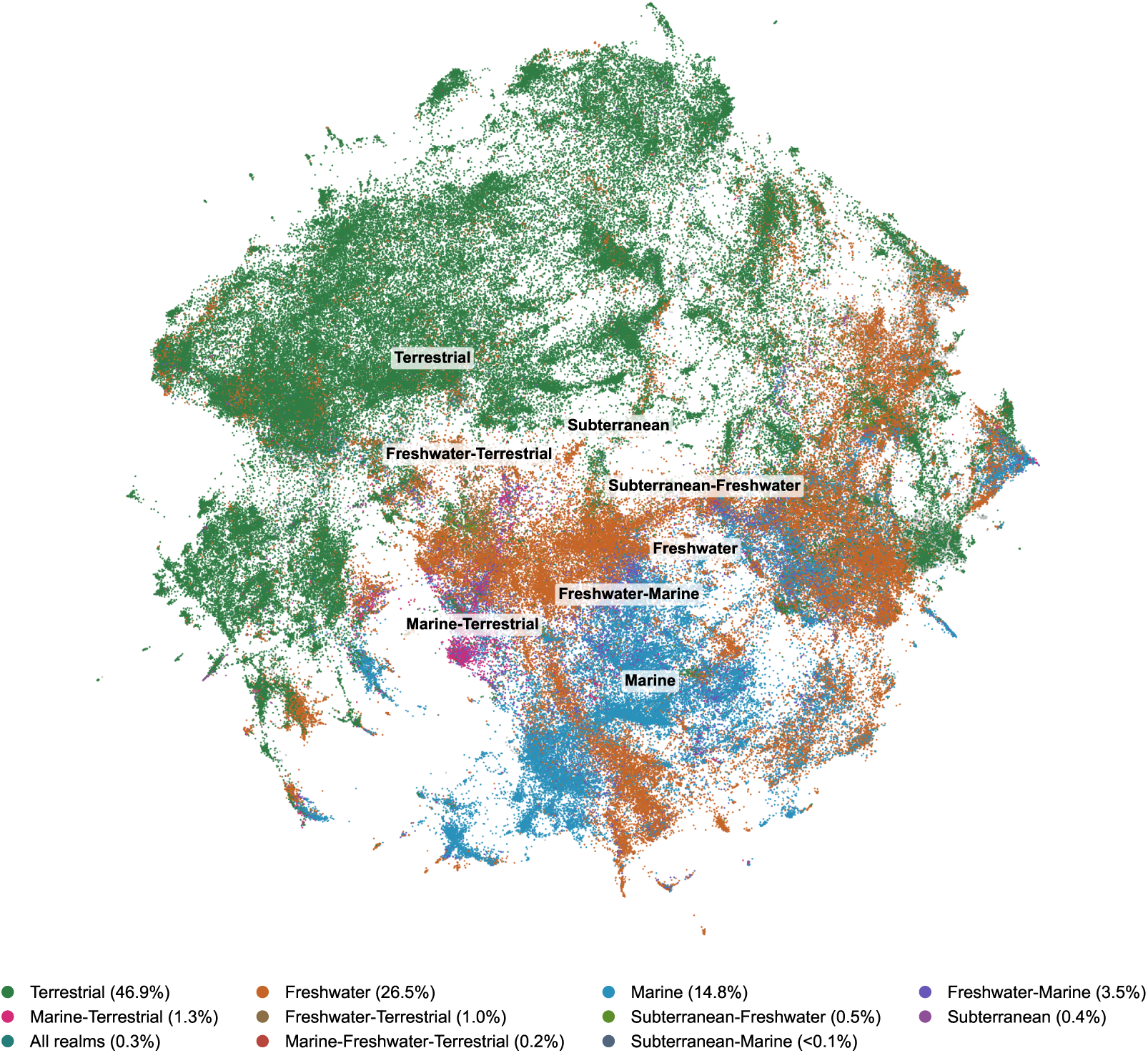
UMAP projection of research on biodiversity loss across ecosystems. Each point represents one of the *n* = 253,081 articles reporting negative biodiversity change, positioned using a two-dimensional UMAP projection of text embeddings of its title and abstract. Colors indicate the ecosystem assigned to the article, and percentages in brackets give the share of articles naming each realm. Articles assigned several ecosystem realms are colored by the most frequent realm across all plotted articles; percentages sum to less than 100% because articles without an assigned realm are not listed. Nearby points represent articles with more similar textual content. Text embeddings were generated using nomic-embed-text-v1.5 and projected using UMAP (n_neighbors = 15, min_dist = 0.1, metric = cosine, random_state = 42).

**Supplementary Figure S8:**
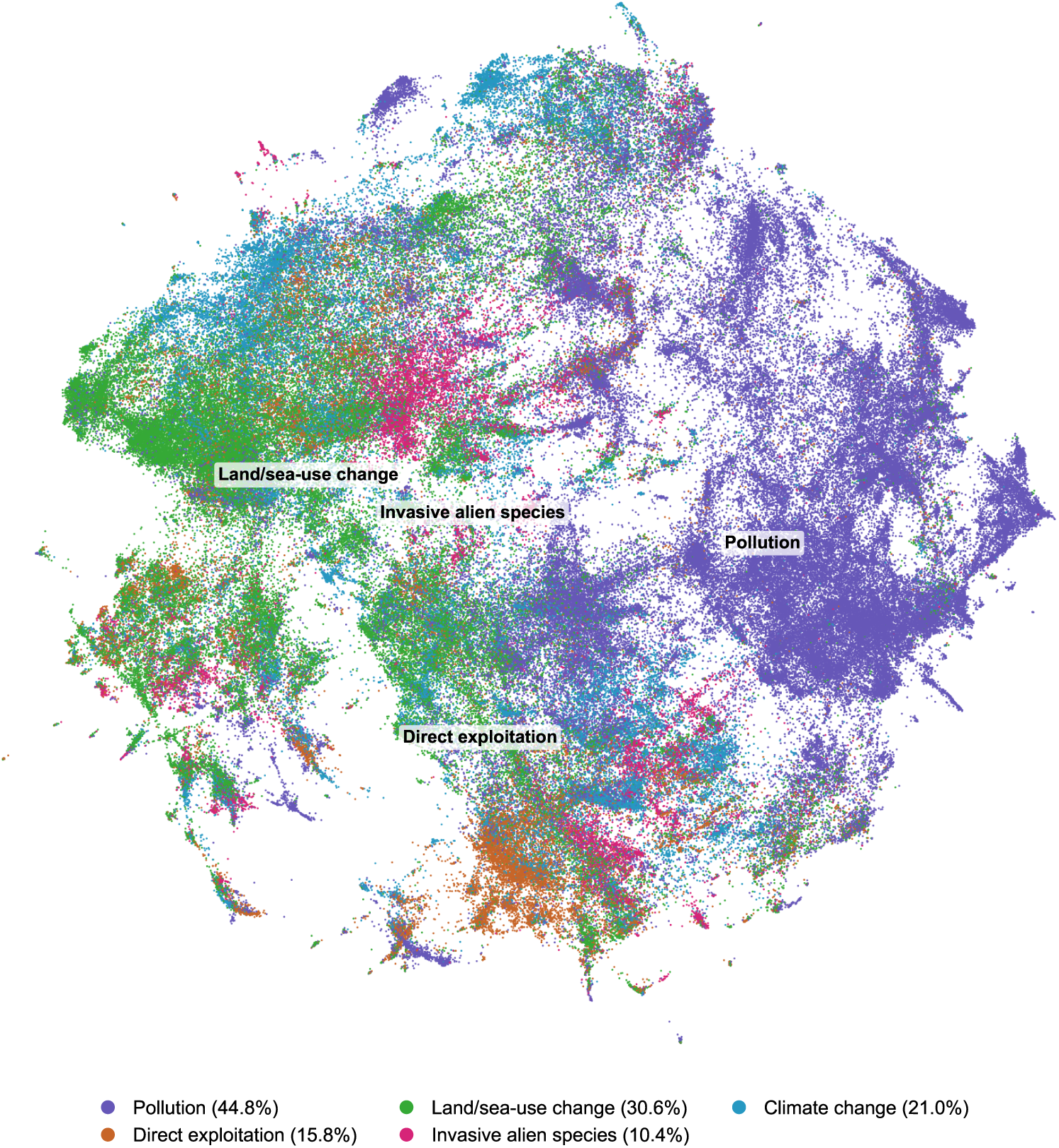
UMAP projection of research on biodiversity loss across direct drivers. Each point represents one of the *n* = 253,081 articles reporting negative biodiversity change, positioned using a two-dimensional UMAP projection of text embeddings of its title and abstract. Colors indicate the IPBES direct driver assigned to the article, and percentages in brackets give the share of articles naming each driver. Articles assigned to several drivers are colored by the driver that is most frequent across all plotted articles (i.e., percentages can sum to more than 100%). Nearby points represent articles with more similar textual content. Text embeddings were generated using nomic-embedtext-v1.5 and projected using UMAP (n_neighbors = 15, min_dist = 0.1, metric = cosine, random_state = 42).

**Supplementary Figure S9:**
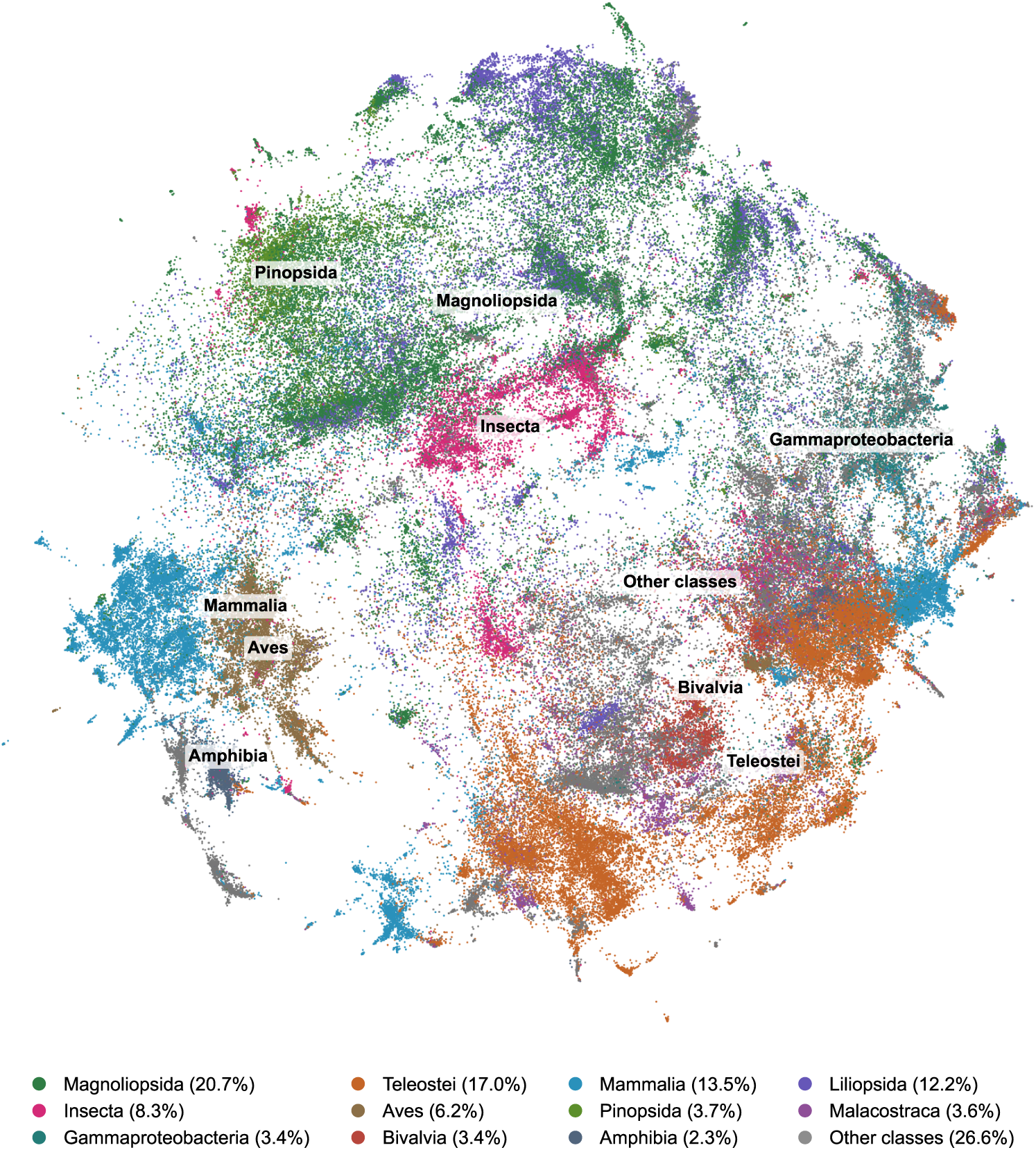
UMAP projection of research on biodiversity loss across taxonomic classes. Each point represents one of the *n* = 163,227 articles reporting negative biodiversity change with at least one taxonomic class label, positioned using a twodimensional UMAP projection of text embeddings of its title and abstract. Colors indicate the 11 most frequent classes; all other classes are pooled as “Other classes”. Percentages in brackets give the share of articles naming each class. Articles naming several classes are colored by the class that is most frequent across all plotted articles (i.e., percentages can sum to more than 100%). Nearby points represent articles with more similar textual content. Text embeddings were generated using nomic-embed-text-v1.5 and projected using UMAP (n_neighbors = 15, min_dist = 0.1, metric = cosine, random_state = 42).

## Supplementary Tables

**Supplementary Table S1:** Changes in research attention across broad IUCN threat categories, 2000–2025. Shares represent the proportion of annual research attention assigned to each threat category, with each article contributing a total weight of one, divided equally among its distinct assigned categories. Changes are expressed in percentage points (pp), calculated as the 2025 share minus the 2000 share. *Other options* and *Unclear* are included in the compositional calculations.

| Broad IUCN threat category | Share 2000 (%) | Share 2025 (%) | Change (pp) |
| --- | --- | --- | --- |
| Residential & commercial development | 1.86 | 5.68 | 3.82 |
| Agriculture & aquaculture | 10.30 | 9.15 | −1.15 |
| Energy production & mining | 1.42 | 1.99 | 0.57 |
| Transport & service corridors | 1.30 | 1.60 | 0.30 |
| Biological resource use | 11.25 | 5.89 | −5.36 |
| Human intrusions & disturbance | 1.27 | 1.02 | −0.25 |
| Natural system modifications | 7.65 | 5.40 | −2.25 |
| Invasive & problematic species | 11.25 | 10.43 | −0.82 |
| Pollution | 40.45 | 32.51 | −7.94 |
| Climate change & severe weather | 4.90 | 19.54 | 14.64 |
| Other options | 1.46 | 1.24 | −0.22 |
| Unclear | 6.88 | 5.54 | −1.34 |

**Supplementary Table S2:**
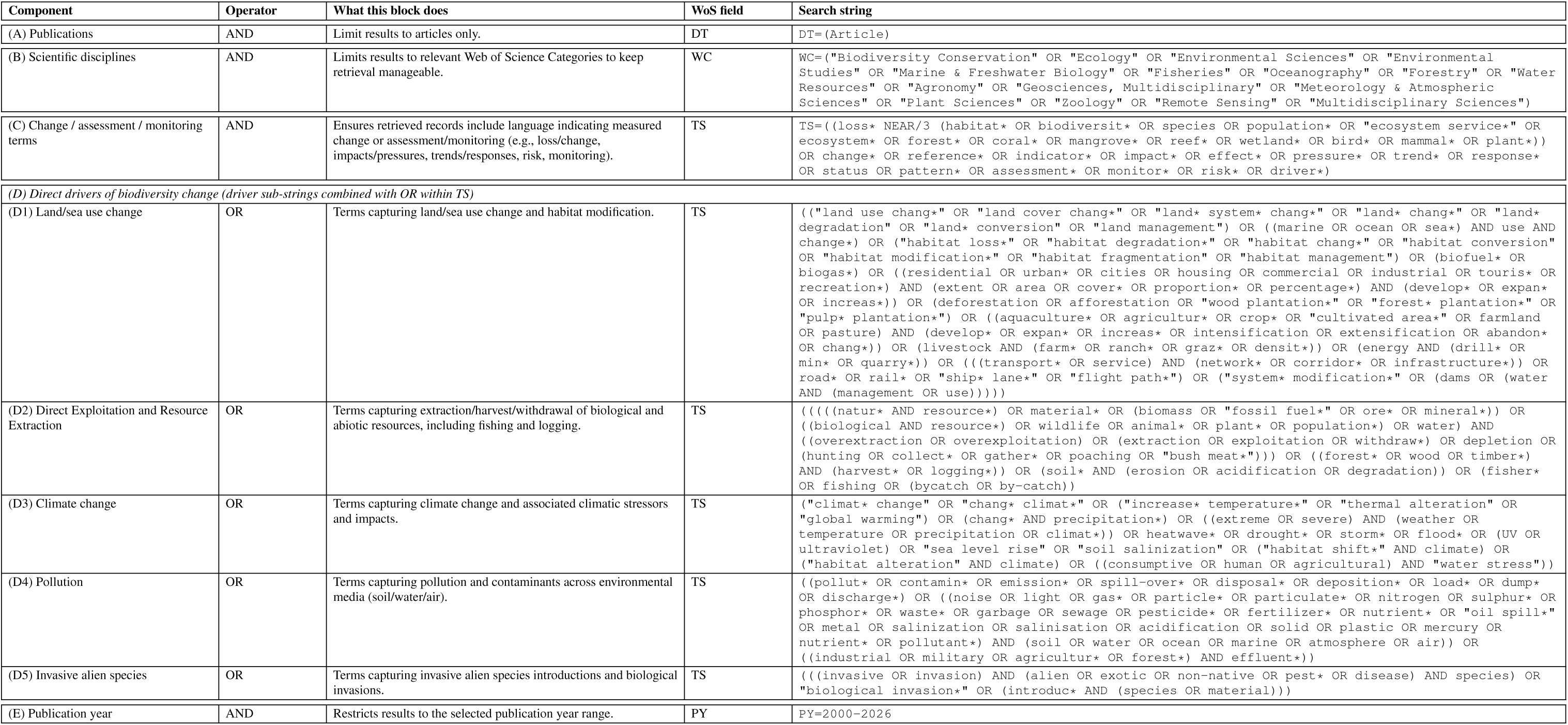
Search string for Web of Science (WoS). Components (A)–(E) are combined using AND. The five sub-strings (D1)–(D5) are taken from [31] and are combined using OR. Component (E) applies the WoS publication-year filter PY=2000-2026; because WoS publication-year filtering does not express a day/month cutoff, records with publication-date metadata after 30 April 2026 were discarded as part of the export.

**Supplementary Table S3:**
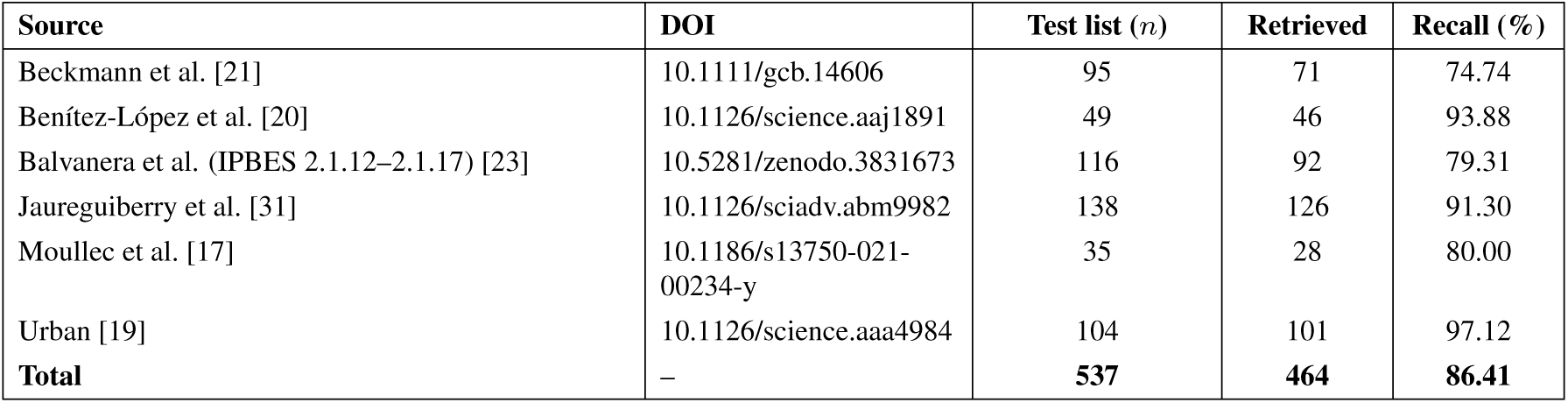
Benchmarking literature retrieval against existing survey articles. For each survey, the test list count is the number of eligible records indexed in WoS and published within the search window (2000 onwards), and recall is the percentage retrieved by the final search string. Overall, there are *n* = 537 articles that we use to benchmark the recall of the search string.

| Source | DOI | Test list ($n$) | Retrieved | Recall (%) |
| --- | --- | --- | --- | --- |
| Beckmann et al. [21] | 10.1111/gcb.14606 | 95 | 71 | 74.74 |
| Benítez-López et al. [20] | 10.1126/science.aaj1891 | 49 | 46 | 93.88 |
| Balvanera et al. (IPBES 2.1.12–2.1.17) [23] | 10.5281/zenodo.3831673 | 116 | 92 | 79.31 |
| Jaureguiberry et al. [31] | 10.1126/sciadv.abm9982 | 138 | 126 | 91.30 |
| Moullec et al. [17] | 10.1186/s13750-021-00234-y | 35 | 28 | 80.00 |
| Urban [19] | 10.1126/science.aaa4984 | 104 | 101 | 97.12 |
| **Total** | – | **537** | **464** | **86.41** |

**Supplementary Table S4:** Distribution of direction of change (Q2) across the 605,199 eligible records. Negative direction constitutes the biodiversity loss dataset analyzed in this study (Supplementary Figure S2); the remaining categories are retained in the eligible set and are released alongside the loss evidence base, though not analyzed in this study.

| Direction of change | Records | % of eligible |
| --- | --- | --- |
| Negative (biodiversity loss) | 253,081 | 41.82% |
| Mixed | 141,267 | 23.34% |
| Positive | 112,684 | 18.62% |
| Unclear | 98,167 | 16.22% |

**Supplementary Table S5:** Validation datasets. Overview of the data used to validate LLM-based eligibility screening and evidence coding.

| **Validation data** | **Eligibility screening** | **Evidence coding** |
| --- | --- | --- |
| Published reference labels | $n = 2,418$ records with published inclusion decisions based on full-text assessments. | $n = 7,261$ articles with published classifications. |
| Manual title-and-abstract labels | $n = 500$ records: 100 sampled from the screening reference data and 400 from the search results. The latter were used for the direct LLM–manual comparison. | $n = 500$ dimension-specific labels (100 per dimension), covering 479 distinct articles. |

**Supplementary Table S6:**
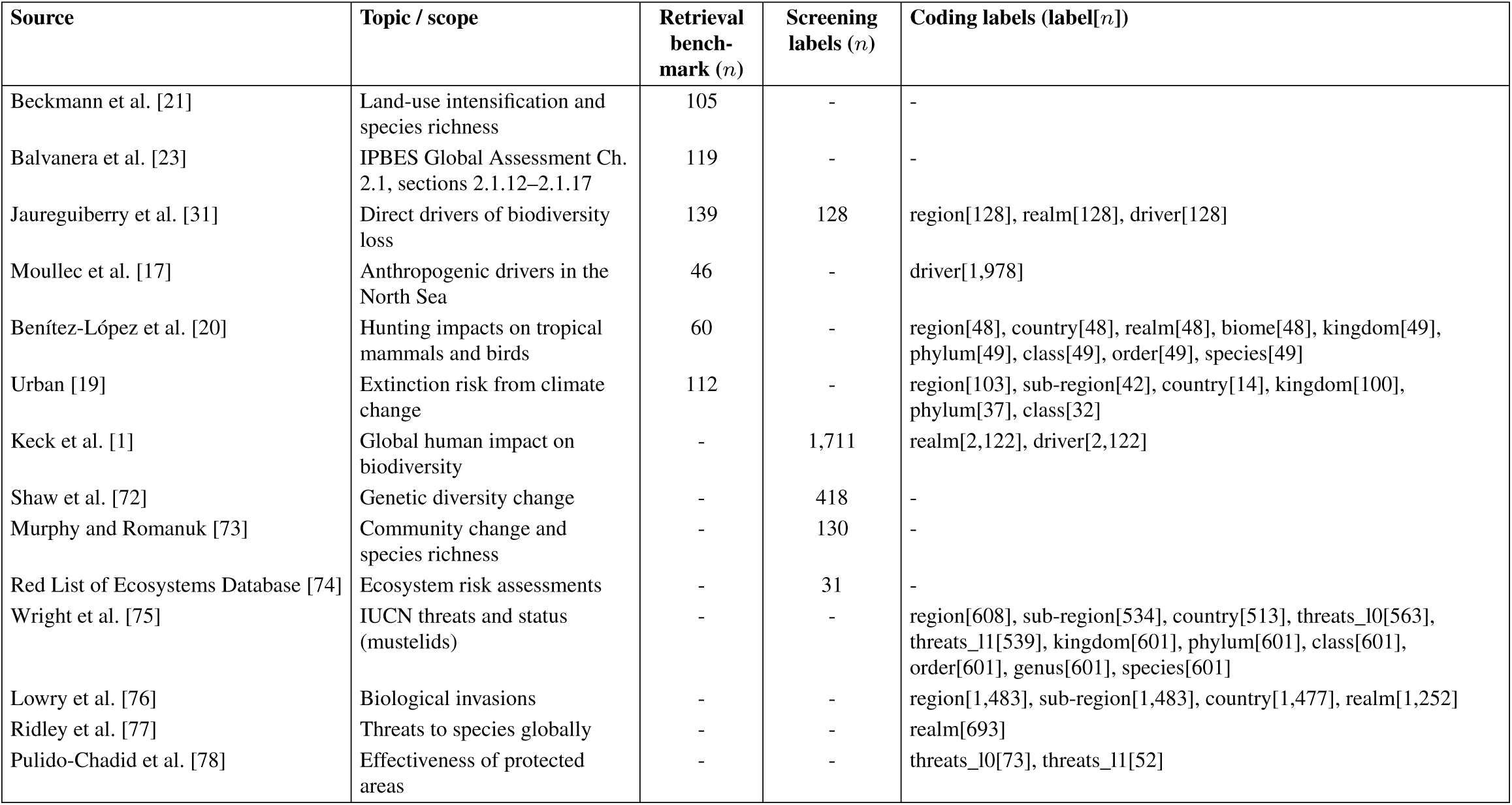
Sources for reference label validation and retrieval benchmarking. Fourteen sources, including one pre-print, contributing to the literature retrieval benchmark or providing published reference labels for eligibility screening and evidence coding. Retrieval benchmark counts refer to the original 581 articles, of which 537 fell within the search period.

**Supplementary Table S7:**
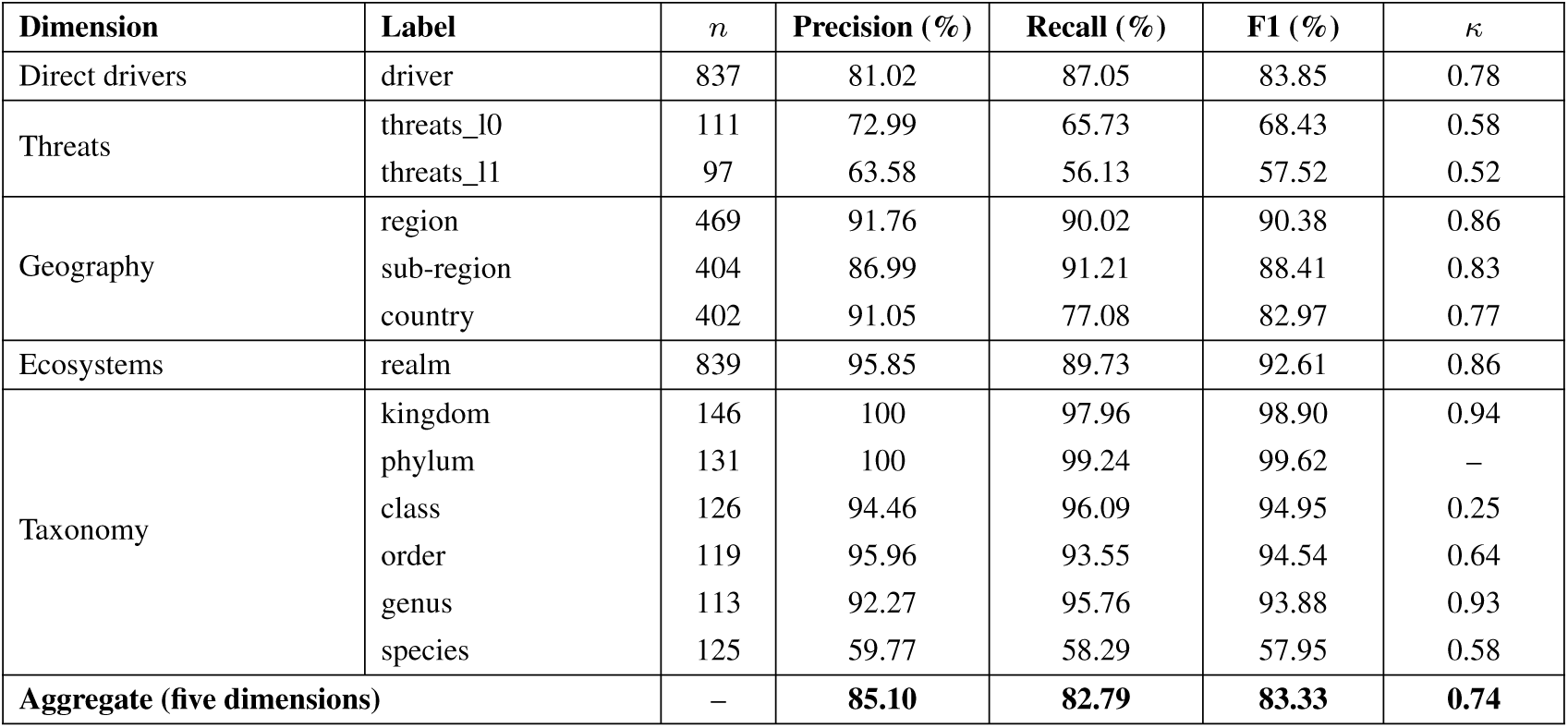
Validation of LLM coding against published reference labels. Field-level performance on held-out test records. *n* denotes the number of scorable records per field after applying the evaluation exclusions. The aggregate represents the macro-average across five dimensions. The phylum test data contained only one observed phylum (*Chordata*); consequently, Cohen’s *κ* was undefined and the high scores should be interpreted cautiously.

**Supplementary Table S8:**
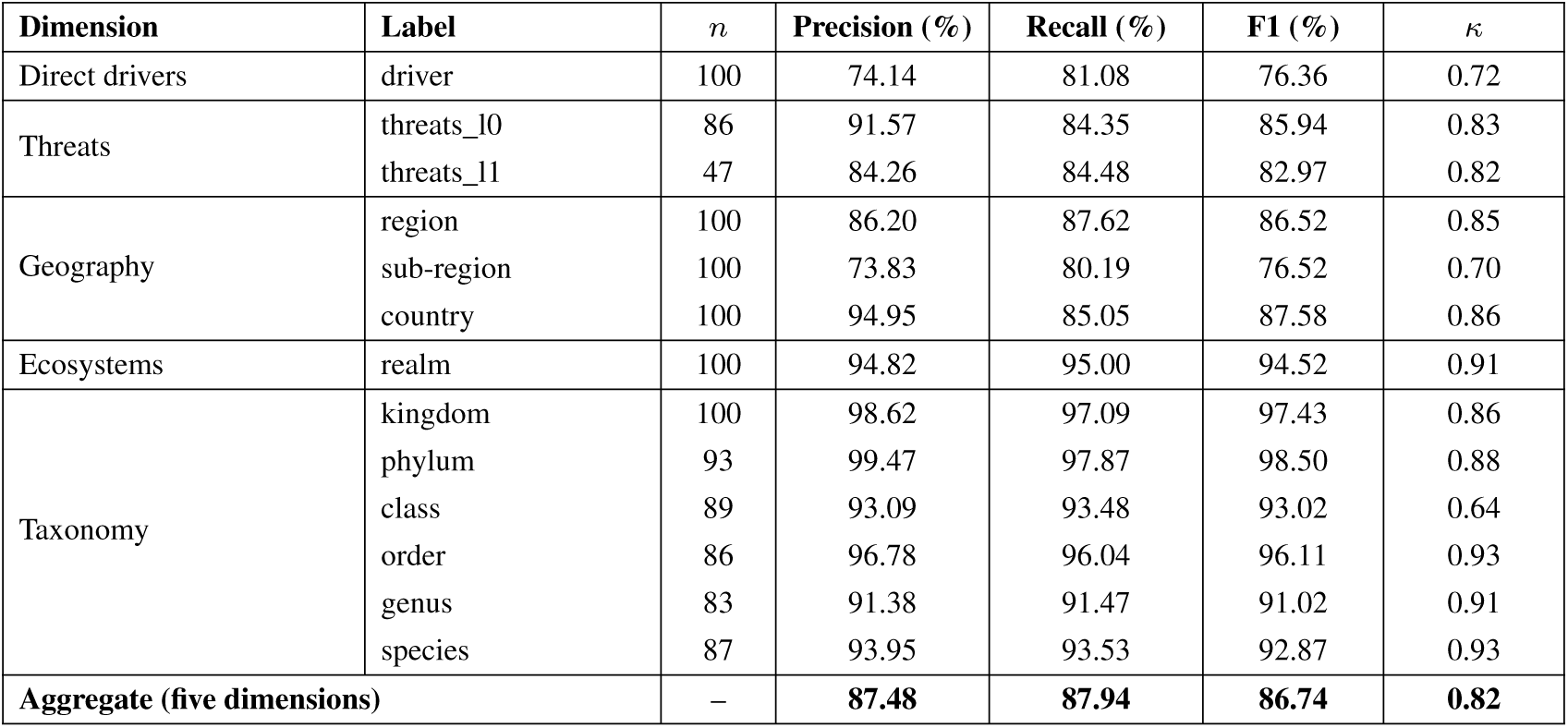
Validation of LLM coding against manual title-and-abstract labels. Field-level results for 500 dimension-specific labels (479 unique articles). *n* denotes scorable records per field after applying the evaluation exclusions. The aggregate follows the same five-dimensional averaging procedure as Supplementary Table S7. Detailed IUCN threat predictions were unavailable for 13 records whose broad-level classification did not trigger the subsequent coding step.

**Supplementary Table S9:** Output schema of the eligibility screening prompt. For each step, the model returned a decision (yes = `1`, no = `0`, unclear = `-1`) and one label, either a single value or a list, with allowed values that depend on the decision. Questions are quoted here as-is from the prompt (`screen.md`).

| Step | Question | Output field | Returned as | Allowed values by decision |
| --- | --- | --- | --- | --- |
| 1 (Q1) | Does the input text report, assess, or clearly imply at least one biodiversity-relevant ecological change in the focal biological system? | biodiversity_types | List (multiple values possible) | Yes (1): one or more of genetic_change, species_change, community_change, ecosystem_change<br>No (0): empty list<br>Unclear (-1): empty list |
| 2 (Q2) | Does the input text report or plausibly indicate the direction of the biodiversity-relevant ecological change? | direction | Single value | Yes (1): one of negative, positive, mixed<br>No (0): none<br>Unclear (-1): unclear |
| 3 (Q3) | Does the input text mention or plausibly imply one or more direct anthropogenic drivers or stressors? | drivers | List (multiple values possible, maximum 8) | Yes (1): one or more free-text names of the drivers identified<br>No (0): empty list<br>Unclear (-1): empty list |
| 4 (Q4) | Does the input text report, assess, attribute, compare, discuss, or clearly imply a link between one or more direct anthropogenic drivers/stressors and the biodiversity-relevant ecological change(s)? | linkage | Single value | Yes (1): one of direct_primary_evidence, modelled, projected, assessed, plausibly_implied, other_positive<br>No (0): none<br>Unclear (-1): unclear |

**Supplementary Table S10:** Prompts, reasoning effort, and output schema for evidence coding. Prompt files are stored in the checklists/prompts folder of the code repository. Effort is the reasoning effort setting. Values per record gives how many values the model could return for a record. Special values (Unclear, Not Applicable, Other Options) are used when a record cannot be assigned to a specific category; the last column explains when each applies.

| Dimension | Prompt file | Effort | Output field | Values per record | Allowed values | Special values |
| --- | --- | --- | --- | --- | --- | --- |
| Direct drivers | `classify_direct_driver` | Low | Labels | One or more | The five IPBES direct drivers [23] | None: every record is assigned at least one driver |
| Threats | `classify_threats` | Medium | Labels | One or more | Categories of the IUCN Threats Classification Scheme [24], Other Options, Unclear | *Other Options*: a direct anthropogenic threat is present but no category fits. *Unclear*: a human impact is described but the text does not support a reliable assignment |
| Geography | `classify_region` | Low | Regions | One or more | IPBES regions [25] (e.g., Africa, Americas), All Regions | *All Regions*: the study is global (not combined with other regions). *Not Applicable*: no region could be assigned |
|  |  |  | Subregions | One or more | IPBES subregions [25] of an assigned region (e.g., Central Asia, South America) | *Not Applicable*: no subregion could be assigned |
|  |  |  | Country codes | One or more | ISO 3166-1 alpha-3 codes, Unclear, Not Applicable | *Not Applicable*: no country could be assigned, for example when only a region is stated. *Unclear*: geography is present but the countries cannot be resolved |
|  |  |  | Locales | One or more | Place names (free text), Unclear, Not Applicable | *Not Applicable*: no locale could be assigned. *Unclear*: geography is present but no locale can be resolved. A placeholder is used if the area is described only by coordinates |
|  |  |  | Coordinates | One per locale | Latitude and longitude for each locale | *Not Applicable*: no coordinates are stated or can be estimated unambiguously |
| Ecosystems, realm | `classify_ecosystem_typology_realm` | High | Labels | One or more | Realms of the Global Ecosystem Typology [26] (e.g., Terrestrial, Freshwater-Marine), Not Applicable, All realms | *Not Applicable*: no real ecological setting. *All realms*: the article is explicitly global across the biosphere. There is no Unclear value |
| Ecosystems, biome | `classify_ecosystem_typology_biome` | High | Labels | One or more | Biomes of the Global Ecosystem Typology [26] within the classified realm(s), Unclear, Not Applicable | *Not Applicable*: no real ecosystem setting. *Unclear*: a setting is present but cannot be mapped to a biome. No biome is recorded if the realm was Not Applicable or All realms, or if the text lacked detail for a biome |
| Taxonomy | `classify_taxa` | High | Taxon rank | One per taxon (several taxa possible) | Kingdom to species, Unclear, Not Applicable | *Unclear*: a taxon is present but cannot be resolved to a name or rank. *Not Applicable*: the text is not clearly about any taxon. Taxa that fit none of the ranks are omitted |
|  |  |  | Canonical name | One per taxon (several taxa possible) | Accepted canonical name in the GBIF Backbone Taxonomy [27], Unclear, Not Applicable | Unclear and Not Applicable are returned together for the rank and the name of a taxon and are not looked up in GBIF. All other names are matched to the GBIF Backbone Taxonomy through the GBIF species-match API |

## Supplementary Methods

### S1 Literature search

The literature search was conducted on the WoS for publications from January 2000 to April 2026, with no language restrictions. The search string was designed to prioritize recall over precision; the search string combines five components using an `AND` operator (see Supplementary Table S2). Component (A) restricts retrieval to journal articles; component (B) limits results to biodiversity-relevant WoS subject categories; component (C) requires terminology indicating measured change, assessment, or monitoring; component (D) requires five driver-specific sub-strings corresponding to the IPBES direct driver categorization (which we adopted from a prior evidence assessment [31]), combined with `OR`; and component (E) applies a publication-year filter.

We assessed the comprehensiveness of our search against an *a priori* benchmark of 581 records drawn from five published sources (i.e., three meta-analyses, an evidence assessment, and a systematic map protocol) focused on studying direct drivers of biodiversity change [17, 19–21, 31] and the IPBES Global Assessment chapters on direct drivers (sections 2.1.12–2.1.17) [23]. The list was assembled by extracting reference lists from each source and excluding grey literature and records not indexed in WoS. Overall, this yielded 581 records from an initial 791 candidates; scientific publications cited by the original authors as evidence were treated as eligible for our evidence map. Of the 581 records, 44 were published before 2000 and therefore lie outside the search window. Of the remaining 537 records, our search string retrieved 464, corresponding to a recall of 86.41%, with source-specific recall ranging from 74.74% to 97.12% (see Supplementary Table S3).

### S2 Eligibility screening

Eligibility screening was performed on titles and abstracts using the four criteria specified in the pre-registered protocol [65]: (Q1) whether biodiversity change was reported, (Q2) the direction of change, (Q3) whether a direct anthropogenic driver was identified, and (Q4) whether the driver was linked to biodiversity change. GPT-5-nano evaluated each criterion using a zero-shot prompt and returned structured Yes, Unclear, or No responses. To prioritize recall, records were retained as eligible unless at least one criterion was explicitly classified as No. The screening prompt and output schema are available in the accompanying code repository.

Of the 2,414,191 records entering eligibility screening, 605,199 (25.1%) were retained as eligible. As specified in the pre-registered protocol [65], eligibility did not require a negative direction of biodiversity change. Of these eligible records, 253,081 (41.82%) reported biodiversity loss and formed the evidence map analyzed in this study. The remaining 352,118 records reported mixed, positive, or unclear directions of change and were retained in the released dataset but are not the focus of this study (Supplementary Table S4). The screening process is documented in the ROSES flow diagram (Supplementary Figure S2), with patterns of exclusion across the four eligibility criteria shown in Supplementary Figure S3.

### S3 Validation

#### S3.1 Validation datasets

We validated LLM-based eligibility screening and evidence coding using four datasets from two complementary sources: (i) published reference labels derived from full-text assessments in previous evidence syntheses, and (ii) manual title-and-abstract annotations. These datasets enabled validation against published reference classifications and direct comparisons between LLM and manual labels on the same records (Supplementary Table S5).

<u>Ad (i):</u> Published reference labels comprised screening inclusion decisions and articlelevel coding classifications from previous evidence syntheses. Screening decisions were based on full-text assessments by the original authors. Source records were matched to their Web of Science entries, and coding labels were harmonized with the geographic, ecosystem, taxonomic, direct-driver, and threat categories used in this study. Published labels served as evaluation references, although the original inclusion criteria and classification conventions did not necessarily coincide with ours. Contributing sources and their respective label coverage are detailed in Supplementary Table S6.

<u>Ad (ii):</u> For the manual comparisons, records were annotated by the authors using titles and abstracts only, without consulting the full texts. For screening, 100 records were drawn from the published screening reference data and 400 records directly from the literature search results; the latter were used for the LLM–manual screening comparison. For evidence coding, 100 records per extracted dimension were sampled from the published coding reference data, with the original reference labels withheld during annotation. The resulting 500 dimension-specific labels represented 479 unique articles and were used for the direct LLM–manual coding comparison.

#### S3.2 Validation of eligibility screening

We validated LLM-based eligibility screening against published reference labels and manual title-and-abstract annotations, as described in Supplementary Methods S3.1.

<u>Ad (i):</u> *Validation against published reference labels.* LLM-based screening was evaluated against published inclusion decisions from previous evidence syntheses. The final screening configuration was selected using development data and evaluated on the heldout test set (*n* = 482), achieving a recall of 90.5%. The published screening reference labels contained only previously included records, so this evaluation focused on recall.

<u>Ad (ii):</u> *Validation against manual title-and-abstract labels.* To assess screening on records drawn directly from the literature search, LLM labels were compared with manual title-and-abstract annotations on a separate sample of 400 records. Relative to the manual labels, LLM screening achieved a precision of 64.5%, recall of 90.1%, and F1 of 75.2%. This comparison complements the published-reference evaluation by including both eligible and ineligible records.

#### S3.3 Validation of evidence coding

We validated LLM-based evidence coding against published reference labels and manual title-and-abstract annotations across 13 classification fields spanning five dimensions: geography, ecosystems, taxa, direct drivers, and threats. The evaluation included both broad (Level 0) and detailed (Level 1) IUCN threat classifications; only Level 0 classifications were retained for full-corpus coding. Coding prompts, configurations, and the complete output schema are available in the accompanying code repository.

Within each classification field, precision, recall, and F1 were calculated using support-weighted averaging. Field-level metrics were then averaged within each dimension and equally across the five dimensions, including both IUCN threat levels. Cohen’s *κ* was aggregated using the same procedure, excluding undefined values. Records with missing, unclear, or inapplicable labels were excluded from scoring, resulting in different sample sizes across fields.

<u>Ad (i):</u> *Validation against published reference labels.* LLM coding was validated against published article-level classifications from previous evidence syntheses and related sources. Prompt configurations were developed and selected using training and development data, with the selected configurations evaluated on held-out test records. Across the five dimensions, the LLM achieved a macro-averaged weighted F1 of 83.33% and Cohen’s *κ* of 0.74 (Supplementary Table S7). Performance varied across classification fields, with lower agreement for detailed IUCN threats (F1 57.52%) and species-level classifications (F1 57.95%) than for broader geographic, ecosystem, and taxonomic categories.

<u>Ad (ii):</u> *Validation against manual title-and-abstract labels.* LLM classifications were also validated against manual annotations of the same titles and abstracts. The comparison comprised 500 dimension-specific labels across five dimensions, representing 479 unique articles. Relative to the manual labels, the LLM achieved a macro-averaged weighted F1 of 86.74% and Cohen’s *κ* of 0.82 (Supplementary Table S8).

**Scope and interpretation.** The five-dimensional validation covered 13 classification fields, while additional outputs, including study attributes, subnational locations, taxonomic family, and biome classifications, were not included in these aggregate performance estimates. Exploratory manual checks of 50 records per field yielded weighted F1 scores of 90.01% for biome classifications and 76.75% for ecosystem functional groups; these smaller assessments were separate from the validation analyses reported above. Detailed IUCN threat classifications and ecosystem functional groups were not retained for fullcorpus coding. Species-level classifications were retained but should be interpreted cautiously given their lower agreement with published reference labels. The complete output field definitions and validation materials are provided in the accompanying code repository.

### S4 Analyses

#### S4.1 Geographic and income-group analyses

Geographic and income-group analyses included 146,630 biodiversity-loss articles published between 2000 and 2025 with study-country assignments resolved to standardized ISO3 country codes and corresponding World Bank classifications, covering 213 countries and territories. Each study country was assigned to the World Bank income group applicable in the article publication year: low-income (LIC), lower-middle-income (LMIC), upper-middle-income (UMIC), or high-income (HIC) [28].

For the country map (Figure 3a), each article was counted once per study country, yielding 165,174 article–country attributions from 146,630 articles. Income-group counts similarly included each article once per represented income group, yielding 151,861 article–income-group attributions. For comparisons of research attention across countries, geographic regions, and income groups, each article instead contributed a total weight of one, divided equally among its study countries and, where applicable, its assigned broad IUCN threat categories. We calculated the composition of research attention within each income group and expressed the difference between HIC and LIC as the HIC-minus-LIC difference in threat share, in percentage points. The corresponding 95% bootstrap confidence intervals were obtained from 1,000 joint publication-level resamples using the 2.5th and 97.5th percentiles of the resulting differences.

#### S4.2 Taxonomic analyses

Taxonomic labels were resolved against the GBIF Backbone Taxonomy [27]. Of the 245,168 biodiversity-loss articles published between 2000 and 2025, 171,791 (70.1%) could be assigned to at least one of five broad taxonomic groups: chordates (Animalia, Chordata), non-chordate animals (Animalia with a resolved phylum other than Chordata), plants (Plantae), fungi (Fungi), and other resolved groups (including Bacteria and Archaea). Articles associated with multiple groups contributed equally to each group.

We compared research attention with the distribution of accepted described species in the GBIF Backbone Taxonomy snapshot of 28 August 2023 [27]. Of 2,614,593 accepted species records, 2,607,214 could be assigned to the broad groups and were included in the benchmark. For comparisons across kingdom, phylum, class, and order, we retained taxonomic paths represented in both the mapped evidence and the benchmark, yielding 113,342 articles in the order-level comparison (Figure 4). Coverage varied across taxonomic groups, with 52% of chordate articles and 74% of articles assigned to other groups represented in the hierarchical comparison, partly reflecting differences in fish and reptile classifications between the two taxonomic sources. The hierarchical and broad-group comparisons thus represent different subsets of the taxonomically resolved literature. Within each rank, articles assigned to multiple taxa contributed equally across their distinct taxonomic paths.

For the regional comparison of chordate research attention (Supplementary Figure S6), each article was counted once in every IPBES region containing a study country. Regional shares were compared with IUCN Red List threatened-species occurrences, defined as species assessed as Critically Endangered, Endangered, or Vulnerable and aggregated across IPBES subregions [29]. The IUCN benchmark covers all comprehensively assessed taxonomic groups rather than chordates specifically and therefore provides broader geographic context for the distribution of research attention.

#### S4.3 Ecosystem analyses

Comparisons of research attention to direct drivers across ecosystems included 215,530 biodiversity loss articles published between 2000 and 2025 that were assigned exclusively to terrestrial (*n* = 114,710), freshwater (*n* = 64,684), or marine (*n* = 36,136) systems. Articles with multiple, transitional, subterranean, or inapplicable ecosystem assignments were excluded. We calculated the distribution of research attention across the five IPBES direct drivers by dividing each article’s total contribution equally among its assigned drivers, so that driver shares summed to 100% within each ecosystem. Annual pollution and direct-exploitation shares were calculated using the same approach (Figure 5; Supplementary Figure S5).

### S5 Prompts and model configuration

We used GPT-5-nano (version `2025-08-07`) through the OpenAI API for both eligibility screening and evidence coding. Requests were submitted in batch mode, and the reasoning effort was set separately for each task. The prompts and output schemas are available in the `checklists/prompts` folder of the project code repository (see Code availability). This section describes the input, configuration, and output schema of each task.

#### S5.1 Eligibility screening

The prompt translates the four eligibility criteria of the protocol [65] into four steps, one for each criterion (`Q1`–`Q4`), and instructs the model to prefer cautious inclusion when evidence is weak. For each step, the model returned a decision (yes = `1`, no = `0`, unclear = `-1`) and a label (see Supplementary Table S9); the decisions determined eligibility as described in Supplementary Section S2.

#### S5.2 Evidence coding

Each classification task used a separate prompt with its own structured output schema (Supplementary Table S10). Each record was submitted in a separate request with taskspecific instructions and its title and abstract as input. For hierarchical threat and ecosystem coding, the parent-level output was also provided as context.

## Notes

https://github.com/forsterkerstin/biodiversity-loss-systematic-map

https://biodiversityliterature.com/

https://osf.io/xg8yq/overview

### Competing Interest Statement

The authors have declared no competing interest.

